# Lipid Driven Inter-leaflet Coupling of Plasma Membrane Order Regulates FcεRI Signaling in Mast Cells

**DOI:** 10.1101/2022.11.24.517890

**Authors:** Gil-Suk Yang, Alice Wagenknecht-Wiesner, Boyu Yin, Pavana Suresh, Erwin London, Barbara A. Baird, Nirmalya Bag

**Author notes:** Program in Cellular and Molecular Medicine, Boston Children’s Hospital, Harvard Medical School, Boston, MA, USA.

## Abstract

Engagement of high affinity immunoglobulin E (IgE) receptor FcεRI with extracellular, multivalent antigen (Ag) stabilizes co-existing ordered and disordered phases in the inner leaflet of the plasma membrane. This optimally controls biochemical interactions between signaling components required for transmembrane (TM) signaling in mast cells. The biophysical organization of the resting inner leaflet is poised to respond appropriately to this extracellular stimulation. The resting inner leaflet is generally less ordered than the outer leaflet, with a lipid composition that does not spontaneously phase separate in model membranes. We proposed that coupling between the two leaflets mediates separation into different phase-like domains in the inner leaflet. To test this hypothesis in live cells, we first established a straightforward approach to evaluate changes in membrane order due to inter-leaflet coupling by measuring inner leaflet diffusion of phase-specific lipid probes with Imaging Fluorescence Correlation Spectroscopy (ImFCS) before and after methyl-α-cyclodextrin (mαCD)-catalyzed exchange of outer leaflet lipids (LEX) with exogenous order- or disorder-promoting phospholipids. We examined the functional impact of LEX by monitoring two Ag-stimulated cellular responses, namely early-stage recruitment of Syk kinase to the inner leaflet and late-stage exocytosis of secretory granules (degranulation). Based on changes in probe diffusion, we observed global increase or decrease of inner leaflet order when outer leaflet is exchanged with order or disorder promoting lipids, respectively, in unstimulated cells. Furthermore, the degree of stimulated Syk recruitment and degranulation correlates with the inner leaflet order of the resting cells, which was varied using LEX. Overall, combined LEX and ImFCS platform provides strong evidence of lipid-based control of stimulated TM signaling in live mast cells. In addition, our functional results imply that resting-state lipid composition and ordering of the outer leaflet sets the ordering of the inner leaflet, likely via interleaflet coupling, and correspondingly modulates TM signaling initiated by antigen-activated IgE-FcεRI.

**STATEMENT OF SIGNIFICANCE:** Coupling between plasma membrane leaflets, which are biochemically and biophysically asymmetric, results in a steady-state membrane organization that is thought to play fundamental roles in cellular functions. Here, we present a straightforward assay built around mαCD-catalyzed lipid exchange (LEX) and Imaging Fluorescence Correlation Spectroscopy (ImFCS) to quantitatively characterize a novel, lipid-driven, interleaflet coupling mechanism and its functional impact in live mast cells. We showed that elevation of outer leaflet lipid order induces ordering throughout the inner leaflet in resting cells. This ordering enhances protein-based reactions during Ag-stimulated FcεRI signaling and consequent cellular response. Overall, we provide a compelling evidence of functional relevance of plasma membrane organizational heterogeneity driven by lipid-based interleaflet coupling.

## INTRODUCTION

Within the plasma membrane, phase-like separation into co-existing ordered and disordered regions critically contributes to transmembrane (TM) signaling stimulated by cell surface immunoreceptors including B cell receptors, T cell receptors, and the high affinity receptor for immunoglobulin E (IgE) FcεRI (1). The biophysical properties of the ordered and disordered regions, respectively, resemble the liquid ordered (Lo) and liquid disordered (Ld) phases observed in model membranes, including giant plasma membrane vesicles (GPMVs) (xyr-4). Chemically, the Lo- and Ld-like phases are enriched with saturated and unsaturated lipids, respectively, and their co-existence in cell membranes is regulated by cholesterol and cortical actomyosin network (5). Lo-like domains in the plasma membrane are known colloquially as “rafts” or “nanodomains” (6).

Diffusion measurements of lipid probes in resting plasma membrane using super-resolution, stimulated emission depletion (STED) microscopy-based fluorescence correlation spectroscopy (FCS) provide reasonable evidence of the presence of nanoscopic (<20 nm) ordered regions (nanodomains) in resting live cells (7, 8). This was further supported by complementary evidence from other high resolution techniques (9, 10). However, these regions are metastable which means that they rapidly disappear from one membrane location and reappear at another location. More recently, our total internal reflection fluorescence microscopy (TIRFM)-based imaging fluorescence correlation spectroscopy (ImFCS) measurements on rat basophilic leukemia (RBL) cells report membrane heterogeneity at diffraction-limited length scale (∼320 nm) (11). This may be interpreted as some membrane areas are enriched with these nanodomains (nanodomain-rich area) compared to other areas (nanodomain-poor area). Again, these areas do not remain at specific locations on the plasma membrane and are likely regulated by the dynamic organization of cortical cytoskeleton (for example, Figure S3 in (11)). This dynamism of phase-like organization in the resting steady-state confers the capacity of plasma membrane to rapidly reorganize into another dynamic steady-state when stimulated. In other words, the resting state of the plasma membrane is poised to undergo reorganization with modulated phase-like separation, which is shown to play decisive roles biological functions.

The functional importance of poised ordered/disordered phase-like separation in resting membrane and its modulation/stabilization after stimulation to facilitate transmembrane signaling is experimentally demonstrated in antigen (Ag)-stimulated IgE-FcεRI signaling system (12). Ag-crosslinking of IgE-FcεRI complex yields receptor nanoclusters (∼70 nm in diameter), which are distributed across the entire plasma membrane (13). Receptor clustering stabilizes the phase-like separation in the inner leaflet of the plasma membrane such that ordered regions coalesce around the receptor clusters in this leaflet (12, 14). Lyn, a src family kinase, phosphorylates tyrosine-based activation motifs (ITAMs) in cytosolic segments of FcεRI (15). Lyn is anchored to the inner leaflet of the plasma membrane via saturated lipid (palmitoyl/myristoyl (PM)) chains and thus preferentially partitions into the ordered phase. By contrast, TM phosphatases which dephosphorylate phospho-ITAMs are excluded from the ordered regions in the vicinity of the crosslinked receptors. This exclusion is driven by steric factors and disorder-preference of the TM phosphatases (12, 16). Thus, the net stabilization of the phase-like separation in the Ag-stimulated condition enhances the probability of Lyn/FcεRI interactions while suppressing phosphatase/FcεRI interactions, thereby tipping the receptor phosphorylation/dephosphorylation balance towards net phosphorylation. The next essential step of mast cell signaling is recruitment of cytosolic Syk kinase to the plasma membrane through its binding to tyrosine phosphorylated FcεRI (17). In this scenario, Syk recruitment depends on tyrosine phosphorylated FcεRI, which depends on coupling with Lyn in ordered membrane regions stabilized by Ag engagement and receptor clustering. Plasma membrane recruitment of Syk kinase and phosphorylation of its substrates yields a cascade of downstream signaling reactions eventually leading to the degranulation, i.e., exocytosis of secretory granules containing β-hexosaminidase and other chemical mediators.

Lipids with unsaturated acyl chains are more abundant than saturated acyl chain lipids in the inner leaflet of the plasma membrane (18). Symmetric lipid bilayers mimicking inner leaflet lipid composition do not phase separate even in the presence of cholesterol (19). Two biophysical models, which may not be mutually exclusive, underlying ordered/disordered phase separation in the inner leaflet of live cell plasma membranes have been recently proposed. First, the Mayor group proposed an inter-leaflet coupling mechanism formed by clustering of dynamic actin filaments and interdigitation of long-chain inner leaflet phosphatidylserine lipids with long-chain outer leaflet phospholipids leading to ordered environment around this cluster (20). The second model is actin-independent interactions of lipid chains where the lipid distribution alone or clustering of lipids or lipid-anchored proteins in the outer leaflet changes and consequently induces inner leaflet phase separation (xyr-23).

Of particular interest is the series of recent experiments in lipid bilayers demonstrating that lipid compositional asymmetry, which is commonly maintained in live cells (24, 25), may drive phase separation in the inner leaflet (xyr-29). Simply, ordered/disordered phase-like separation in outer leaflet lipids can induce phase-like separation in the inner leaflet. On the other hand, under some conditions, signal transduction is associated with a loss of lipid asymmetry (30), and a loss of lipid asymmetry may induce ordered domain formation in the inner leaflet, as shown in model lipid bilayers (28) and cell-derived GPMVs (29) systems. While GPMVs are a very useful cell-derived model system to study membrane organization, they lack cortical actin cytoskeleton and suffer from loss of compositional asymmetry to some degree (29). A systematic live cell study on this biophysical coupling between the leaflets and its physiological implications therefore remains highly desirable.

To evaluate the degree of inter-leaflet coupling resulting from lipid compositional asymmetry, we modify the outer leaflet lipid composition by methyl alpha cyclodextrin (mαCD)-catalyzed lipid exchange (LEX) (31, 32) and monitor the corresponding changes of diffusion properties in the inner leaflet (Fig. 1). In this LEX process, a selected exogenous lipid (Lipid_ex_) is complexed with a molar excess of mαCD in solution. This is added to replace endogenous lipids which form complexes with the excess (uncomplexed) mαCD while the Lipid_ex_ (initially complexed with mαCD) is transferred to the outer leaflet. Further investigations (32) show that the Lipid_ex_ remains in the outer leaflet for sufficiently long time (∼2 hr) for our biophysical and functional measurements. The efficiency of the LEX method has so far been characterized in several cell types (reviewed in (33)). It has been shown that ∼50-100% of endogenous phospholipids in the plasma membrane outer leaflet (and likely that of membrane compartments rapidly cycling to and from the plasma membrane (34)) may be exchanged by this method without significantly compromising cell health as judged from no change in cell morphology and membrane integrity (31, 32). These results motivated us to use LEX to easily change outer leaflet lipid composition and test its inter-leaflet effects in live cells.

**Figure 1.**
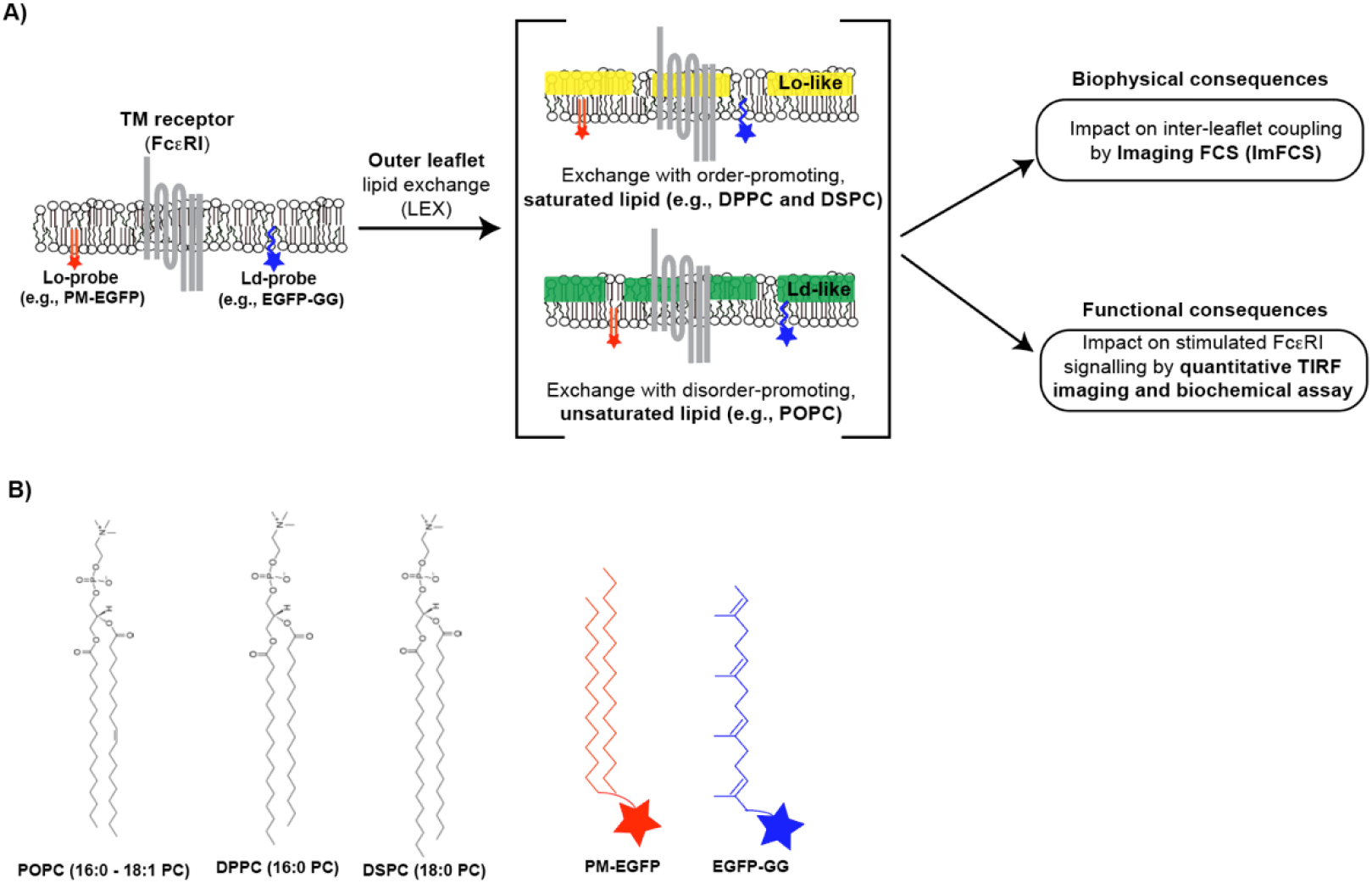
Schematics of experimental strategy to evaluate inter-leaflet coupling of cellular plasma membrane. A) Experimental strategy to perform mαCD-catalyzed outer leaflet lipid exchange (LEX) with exogenous lipid (Lipid_ex_) and evaluate of its biophysical and functional consequences. B) Chemical structures of Lipid_ex_ and inner leaflet lipid probes used in this study. The lipid structures are takes from Avanti Polar Lipids website.

Modulation of phase-like organization in the inner leaflet due to the LEX-based perturbations in the outer leaflet can be evaluated from the corresponding changes in the inner leaflet diffusion. To detect subtle changes in phase-like organization in the plasma membrane, we recently developed a highly robust experimental strategy based around precise (standard error of mean <1%) diffusion coefficient (*D*) measurements of passive (non-functional) lipid probes by Imaging Fluorescence Correlation Spectroscopy (ImFCS) (11, 12). ImFCS measures several thousands of *D* values from multiple cells at a given condition. The entire pool of *D* values is used to compare the difference of diffusion coefficients across experimental conditions (e.g., Ag-stimulated vs unstimulated conditions as in (12)) while the consistency of *D* values across cells is confirmed by bootstrapping (12). We previously showed that diffusion of an order-preferring lipid probe, namely enhanced green fluorescent protein (EGFP) tagged palmitoyl/myristoyl (PM) lipid chains (PM-EGFP), decreases by 8% while that of a disorder-preferring lipid probe, namely EGFP-tagged geranylgeranyl lipid chain (EGFP-GG), increases by 7% upon clustering and stimulation of IgE-FcεRI with multivalent antigen (Ag), DNP-BSA (12). These contrasting diffusion changes of order- and disorder-preferring probes after receptor stimulation reflect relative stabilization of phase-like separation of order and disordered regions in the inner leaflet, to which PM-EGFP and EGFP-GG are anchored. Such stabilization signifies more distinctive ordered and disordered regions causing enhanced partitioning of lipid probes in their preferred regions. Since ordered regions are more viscous than disordered regions, the diffusion of order-preferring PM-EGFP decreases while that of disordered-preferring EGFP-GG increases. Likewise, Shelby et al. used super-resolution localization microscopy to show nanoscale colocalization between PM probe and Ag-clustered IgE-FcεRI, whereas the GG probe did not show any detectable colocalization under the same conditions (14). Collectively, ImFCS and super-resolution imaging show stabilization of ordered regions around the clustered IgE-FcεRI since PM and GG probes do not undergo any protein-based interaction and they respond only to changes in the lipid-based interactions.

Here we use the same ImFCS strategy to investigate the modulation of phase-like organization in the inner leaflet due to LEX of outer leaflet lipids in resting RBL cells. We evaluate the diffusion properties of inner leaflet probes PM-EGFP and EGFP-GG when outer leaflet lipid composition is exchanged with order-promoting saturated Lipid_ex_ including 1,2-dipalmitoyl-sn-glycero-3-phosphocholine (DPPC) and 1,2-distearoyl-sn-glycero-3-phosphocholine (DSPC); or disorder-promoting unsaturated Lipid_ex_, 1-palmitoyl-2-oleoyl-glycero-3-phosphocholine (POPC) (35). We observe that high concentrations of these order- and disorder-promoting Lipid_ex_ species in the outer leaflet respectively decrease and increase diffusion coefficients for both PM-EGFP and EGFP-GG. This correlation reveals that the two leaflets of the plasma membrane are biophysically coupled and the outer leaflet lipid substitution by LEX method induces global changes in the inner leaflet. Since plasma membrane heterogeneity in the resting state is poised to respond Ag-stimulation, it is essential to test the effects of LEX in resting conditions on the functional consequences in stimulated FcεRI signaling. We extend this investigation further into the functional consequences of this lipid-based inter-leaflet coupling. We monitor the extent of stimulated Syk recruitment (early signaling event) and degranulation (ultimate cellular event) in response to the changes in the outer leaflet lipid composition made in resting cells. Our results show clear correlation between LEX-induced changes in the inner leaflet order in the resting state and these two stimulated functional events. Overall, this study provides novel evidence of functional, inter-leaflet coupling in the plasma membranes of live cells.

## MATERIALS AND METHODS

### Reagents

Minimum essential medium (MEM), Dulbecco’s Modified Eagle Medium (DMEM), Opti-MEM, Trypsin-EDTA (0.01%), and gentamicin sulfate were purchased from Life Technologies (Carlsbad, CA). Fetal Bovine Serum (FBS) was purchased from Atlanta Biologicals (Atlanta, GA). Alexa Fluor 488 (AF488) NHS ester (Invitrogen) was used to fluorescently label monoclonal anti-DNP (2,4-dinitrophenyl) immunoglobulin E (IgE) yielding AF488-IgE as described previously (27). The multivalent antigen (Ag), DNP-BSA, was prepared by conjugating DNP sulfonate (Sigma-Aldrich) to bovine serum albumin (BSA) (36). Methyl alpha cyclodextrin (mαCD) was purchased from AraChem (Tilburg, Netherlands). The stock solution of mαCD was prepared in Dulbecco′s Phosphate Buffered Saline (DPBS) buffer and stored at 4°C. Phorbol 12,13-dibutyrate (PDB) was obtained from Sigma-Aldrich (St. Louis, MO). Stock solution of PDB was prepared in DMSO and stored at -80°C. All lipids were purchased from Avanti Polar Lipids (Alabaster, AL) in chloroform stock solution.

### Cell culture, chemical transfection, sensitization, and stimulation

Both wildtype and Syk-negative RBL cells derived from the parental line (37) were grown and maintained in culture medium containing 80% MEM, 20% FBS, and 10 mg/L gentamicin sulfate at 37°C and 5% (v/v) CO_2_ environment. Both cell lines were transfected with FuGENE HD transfection kit (Promega). The details of transfection protocol are described elsewhere (12). The chemically transfected plasmids used in this study encode the following proteins: PM-EGFP (38), EGFP-GG (38), YFP-Syk (39).

All ImFCS experiments were done in resting cells while the functional assays (YFP-Syk recruitment and degranulation) were done on stimulated cells. We describe the sensitization and stimulation conditions for the functional assays below.

For YFP-Syk recruitment assay, YFP-Syk expressing cells were incubated overnight in 35 mm glass bottomed dish (MatTek; Ashland, MA) with 0.5 μg/mL anti-DNP IgE at 37ºC and 5% (v/v) CO_2_ environment followed by LEX in room temperature for 1hr (details of LEX protocol are given in next sections). After LEX, these cells were stimulated with 0.5 μg/mL DNP-BSA at room temperature, and time-lapse TIRFM imaging (see below for the imaging set-up) was done right after addition.

For degranulation experiments, cells were plated in a 96-well plate. The conditions of IgE sensitization and LEX were same as above (for YFP-Syk recruitment assay). These cells were stimulated with 0.01 μg/mL DNP-BSA for 1 hr at 37ºC. Degranulation of the cells was then measured by UV-Vis spectroscopy.

### Preparation of LEX working solution

This protocol and LEX (next section) are adopted from previously published reports (31, 32, 40). Appropriate volume of lipid (in chloroform) in a 5 mL glass vial was first dried under nitrogen gas stream, followed by high vacuum for 1 hour. For stock solution, the dried lipid film was then hydrated in 40 mM mαCD solution at 70°C for 5 min followed by at 55°C for 30 min. The lipid exchange solution thus prepared was kept overnight at room temperature. The working solution of mαCD (40 mM) is prepared by diluting the stock solution (prepared in DPBS buffer) in serum-free DMEM right before addition to the lipid film. The final concentration of the lipids used were 3 mM POPC, 1.5 mM DPPC, and 1.5 mM DSPC.

### Outer leaflet lipid exchange (LEX) in RBL cells

RBL cells grown unto 80-90% confluency on a 35 mm glass bottomed dish (for ImFCS and YFP-Syk recruitment assay) or a 96-well plate (for degranulation assay) were washed twice with buffered saline solution (BSS: 135 mM NaCl, 5.0 mM KCl, 1.8 mM CaCl_2_, 1.0 mM MgCl_2_, 5.6 mM glucose, and 20 mM HEPES; pH 7.4) and then incubated at room temperature for 45 min in fresh BSS. After this, for cells in glass-bottomed dish, BSS was removed from the dish and 100 μL of the LEX working solution containing selected Lipid_ex_ was added on the glass surface of the dish. In this step for 96-well plate, BSS was removed and 100 μL of the LEX solution is added to each well. The dish or the 96-well plate was then incubated in the room temperature for 1 hour while gently shaken. After this, the LEX solution was discarded, and the cells were washed twice with BSS buffer. The cells in the dish were then imaged in fresh BSS buffer. For the control (unexchanged) sample, the cells were subject to same preparation steps as above except for 100 μL of serum-free DMEM was added instead of LEX working solution. All imaging experiments, unless otherwise indicated, were performed within 45 min after the LEX incubation. The degree of lipid exchange for various Lipid_ex_ is monitored by measuring relative reduction of sphingomyelin in whole cell lysates by high-performance thin layer chromatography (HP-TLC) following the protocol described elsewhere (31, 32).

### Degranulation assay

Stimulated degranulation was measured using standard β-hexosaminidase assay. The procedure is described elsewhere (41).

### TIRFM imaging and ImFCS measurements

The instrumental set up for time-lapse TIRFM imaging was described elsewhere (6, 8, 25). TIRFM images of IgE-sensitized, YFP-Syk expressing cells were taken at every 1 min after addition of DNP-BSA. The YFP-Syk puncta analysis (number of puncta per unit area of background-corrected cell images) was done using built-in functions in FIJI/ImageJ (42).

The details of ImFCS data acquisition, analysis, and synthesis of bootstrapped cumulative distribution functions of diffusion coefficient was described previously (12). The ImFCS data acquisition protocol was carried out as described elsewhere (43, 44). ImFCS analysis (45) to determine pixel-by-pixel diffusion coefficient values was done using as FIJI/ImageJ plug-in (Imaging_FCS 1.491; available at http://www.dbs.nus.edu.sg/lab/BFL/imfcs_image_j_plugin.html). All imaging measurements were done at room temperature.

### Quasi-TIRFM imaging

The same TIRFM set up described above was used to do the quasi-TIRFM imaging. The incident angle was manually set at less than critical angle such that the excitation beam is not internally reflected but rather passes through a cross-section of the sample. In this manner, we imaged sample planes away from the surface.

### FRAP measurements

The instrumental set up as well as FRAP data acquisition and fitting criteria were described elsewhere (12).

## RESULTS

The substitution of outer leaflet lipids by selected exogenous lipids (Lipid_ex_) in complexes with mαCD (mαCD-Lipid_ex_), dubbed LEX, is an efficient method to modify outer leaflet composition in a highly controlled manner (31). In this study we extend LEX, which has been primarily used to explore inter-leaflet coupling in model systems, to investigate the same coupling process in live cells and the accompanying effects on cellular signaling. We use RBL mast cells as representative mammalian cell line for which a large body of biophysical, biochemical, and cell biological information is available. For example, phospholipid composition of RBL plasma membrane (46, 47) and biophysical properties of live cell membranes (48, 49) as well as RBL-derived giant plasma membrane vesicles (GPMVs) (2, 50, 51) have been reported. Likewise, mast cell signaling cascade mediated by IgE-FcεRI and the importance of membrane biophysical properties in the earliest stage of this process are well established (12, 14, xyr-54).

### LEX efficiently replaces outer leaflet lipids of RBL plasma membrane

We first characterized the compatibility of LEX for RBL cells. For this, Lipid_ex_ (POPC (3 mM) or DPPC (1.5 mM) or DSPC (1.5 mM)) is complexed with an excess of mαCD (40 mM) such that free mαCD may extract lipids from the plasma membrane while the mαCD-Lipid_ex_ complex delivers the Lipid_ex_ to the plasma membrane (33). The choice of Lipid_ex_ content can be guided by previous report (32).

The plasma membrane pool of sphingomyelin (SM), which is about 70% of whole cell SM content, is almost exclusively present in the outer leaflet (18). As shown previously for other cell types (32, 33, 40), the amount of SM remaining with the cells after LEX is a good indicator of the lipid exchange efficiency. We used high-performance thin layer chromatography (HP-TLC) to determine the level of SM (normalized against total protein concentration) in whole cell lysates in unexchanged (control) and various LEX conditions on RBL cells. Results are shown in Fig. S1. We found that the SM content is reduced significantly under LEX conditions. Specifically, the amounts of residual SM are 23%, 39%, and 48% compared to control cells after exchange of POPC, DPPC, and DSPC, respectively, under our conditions. Correspondingly, the trend of LEX efficiency is: POPC > DPPC > DSPC. The observed amounts of residual SM in RBL cells are comparable to those of Chinese Hamster Ovary cells (40) for these Lipid_ex_ species. As a positive control, we used brain SM (bSM; 1.5 mM) as Lipid_ex_ and observed a marked increase of total SM as expected. Overall, LEX in RBL cells appears to be as efficient as for other cell types tested previously.

### Diffusion in plasma membrane is perturbed by LEX and recovers over prolonged incubation

We started our investigations with ImFCS measurements on Alexa fluor 488 labelled IgE-FcεRI complex (AF488-IgE-FcεRI) before (control) and after POPC-LEX (Fig. S2). RBL cells were labelled with AF488-IgE before LEX. The cells were washed after LEX with BSS, and diffusion of AF488-IgE-FcεRI was measured as a function of time. The average diffusion coefficient (*D*_*av*_) of this transmembrane probe is two times the value for POPC-LEX (*D*_*av*_ = 0.30 ± 0.002 μm^2^/s; mean ± SEM) cells than for control unexchanged cells (*D*_*av*_ = 0.15 ± 0.001 μm^2^/s), and it remains steady until ∼45 min post LEX. Then, the *D*_*av*_ of this probe decreases monotonically over time suggesting recovery of original membrane viscosity. The *D*_*av*_ value is close to that of control cells after a recovery period of ∼2 hr (*D*_*av*_ = 0.19 ± 0.001 μm^2^/s). Based on these observations, we decided to conduct all ImFCS experiments within 45 min after LEX to measure the effects of the perturbation of lipid composition before spontaneous recovery and restoration of normal membrane diffusion occur.

### Changes in the inner leaflet diffusion correlate with ordering and disordering of the outer leaflet

After establishing the measurement criteria, we tested the effects of LEX on the diffusion properties of the inner leaflet to quantitatively evaluate the degree of inner-leaflet coupling. We used Lipid_ex_ species that are either order-promoting (DPPC and DSPC) or disorder-promoting (POPC). To evaluate the inter-leaflet impact of these substantial outer-leaflet changes (Fig. S1), we measure the diffusion of inner leaflet lipid probes PM-EGFP (order-preferring) and EGFP-GG (disorder-preferring) (38) by ImFCS. We used these probes previously to demonstrate local stabilization of ordering around Ag-clustered IgE-FcεRI (12, 14, 38).

Fig. 2A shows bootstrapped cumulative distribution function (CDF) of *D* values of PM-EGFP under various conditions. The shifts in the CDFs provide easy visual distinction of diffusion changes between two conditions (12): a left shift means slower diffusion and vice versa. The diffusion of PM-EGFP in POPC-LEX cells is markedly faster than for control cells indicating strong inter-leaflet interaction (Fig. 2A). The *D*_*av*_ values for PM-EGFP in control and POPC-LEX cells are 0.63 ± 0.004 and 0.90 ± 0.005 μm^2^/s, respectively (Fig. 2A, top). In contrast, the *D*_*av*_ of PM-EGFP is considerably slower in DPPC-LEX (Fig. 2A, middle) and DSPC-LEX (Fig. 2A, bottom) cells than in the control cells measured in the same experiment. The *D*_*av*_ values are 0.62 ± 0.002 and 0.39 ± 0.001 μm^2^/s respectively for corresponding control and DPPC-LEX cells (Table 1). Likewise, the *D*_*av*_ values were 0.67 ± 0.003 and 0.29 ± 0.002 μm^2^/s, respectively, for corresponding control and DSPC-LEX cells (Table 1). As seen here, measurements on the same control cells may vary a bit on different days. Therefore, we determined the ratio of *D*_*av*_ of LEX to control cells as *D*_Lex_/*D*_Ctrl_ parameter to make a more direct and objective comparison across different Lipid_ex_ (Fig. 2B). This parameter is equal to, greater, or lesser than 1 for no change, increase, or decrease, respectively, of *D*_*av*_ due to LEX. The *D*_Lex_/*D*_Ctrl_ values of PM-EGFP for POPC-, DPPC-, and DSPC-LEX are 1.43, 0.63, 0.44 respectively (Fig. 2B and Table 1).

**Table 1:** *D*_*av*_ and *D*_Lex_/*D*_Ctrl_ values for all probes in control and Lipid_ex_-LEX treated cells.

| Probe | Condition | $D_{av}$ [$\mu\text{m}^2/\text{s}$]<sup>a</sup> | $D_{Lex}/D_{Ctrl}$<sup>b</sup> |
| --- | --- | --- | --- |
| PM-EGFP | Control | $0.63 \pm 0.004$ | 1 |
| | POPC-LEX | $0.90 \pm 0.005$ | 1.43 (1.36 – 1.50) |
| | Control | $0.62 \pm 0.002$ | 1 |
| | DPPC-LEX | $0.39 \pm 0.001$ | 0.63 (0.61 – 0.65) |
| | Control | $0.67 \pm 0.003$ | 1 |
| | DSPC-LEX | $0.29 \pm 0.002$ | 0.44 (0.42 – 0.46) |
| EGFP-GG | Control | $0.66 \pm 0.005$ | 1 |
| | POPC-LEX | $0.88 \pm 0.006$ | 1.35 (1.27 – 1.42) |
| | Control | $0.66 \pm 0.003$ | 1 |
| | DPPC-LEX | $0.45 \pm 0.002$ | 0.68 (0.66 – 0.71) |
| | Control | $0.69 \pm 0.003$ | 1 |
| | DSPC-LEX | $0.31 \pm 0.001$ | 0.45 (0.43 – 0.47) |
| FAST-DiO | Control | $0.79 \pm 0.006$ | 1 |
| | POPC-LEX | $0.78 \pm 0.003$ | 0.99 (0.94 – 1.03) |
| | DPPC-LEX | $0.33 \pm 0.002$ | 0.42 (0.40 – 0.45) |
| | DSPC-LEX | $0.22 \pm 0.002$ | 0.28 (0.26 – 0.30) |
| YFP-GL-GPI | Control | $0.38 \pm 0.002$ | 1 |
|  | POPC-LEX | N.D. | N.D. |
| | DPPC-LEX | $0.29 \pm 0.002$ | 0.77 (0.74 – 0.81) |
| | DSPC-LEX | $0.19 \pm 0.001$ | 0.49 (0.47 – 0.51) |
| AF488-CTxB-GM1 | Control | $0.15 \pm 0.001$ | 1 |
| | POPC-LEX | $0.24 \pm 0.001$ | 1.60 (1.51-1.71) |

<sup>a</sup> $D_{av}$ values are given as mean ± SEM
<sup>b</sup> Mean values of $D_{Lex}/D_{Ctrl}$ are provided. In the parenthesis, the range of values of this parameter is given.

**Figure 2.**
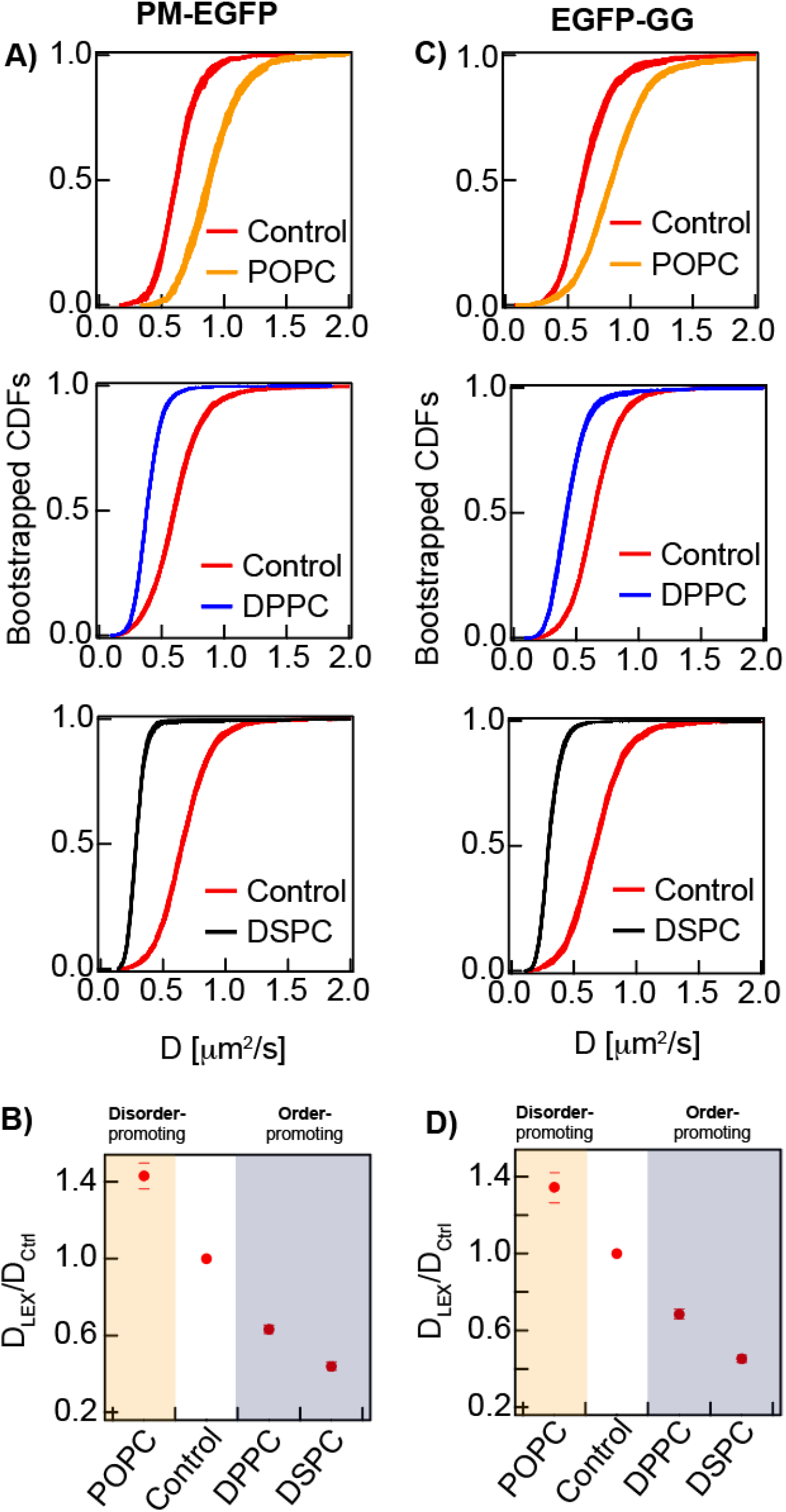
Inter-leaflet coupling of LEX in outer leaflet are sensed by inner leaflet lipid probes. (A-B) Lo-preferring PM-EGFP and (C-D) Ld-preferring EGFP-GG. (A and C) The modulation of probes’ diffusion due to LEX for the indicated types of exogeneous lipids is shown as bootstrapped CDFs of *D* at different control and LEX conditions. (B and D) The relative change of *D* is shown as the *D*_Lex_/*D*_Ctrl_ ratio. The error bars in these plots depict the entire range of values of this parameter.

We conducted the same set of experiments for the disorder-preferring probe EGFP-GG. The CDFs of *D* values for this probe at various LEX conditions are shown in Fig. 2C. The trends of LEX induced diffusion change of EGFP-GG are same as those of PM-EGFP (Fig. 2A): POPC-LEX increases diffusion of both probes while DPPC- and DSPC-LEX decrease their respective diffusion. This similarity in the trends of these two inner leaflet probes may be seen quantitatively when their respective *D*_Lex_/*D*_Ctrl_ values are compared for each Lipid_ex_ (Figs. 2B and D, Table 1). The *D*_Lex_/*D*_Ctrl_ values of EGFP-GG are 1.35, 0.68, and 0.45 for POPC-, DPPC-, and DSPC-LEX, respectively, which are comparable to those of PM-EGFP. These results on the behavior of two probes having orthogonal phase preference are thus consistent with the view that exchange of outer leaflet lipids has inter-leaflet impact in which more disordered outer leaflet globally causes the inner leaflet to be more disordered and vice versa (see Discussion).

### Diffusion of outer leaflet lipid probes is modulated by LEX

LEX mediated inter-leaflet coupling leads to global changes in the inner leaflet diffusion. This is an interesting difference from the effects of nanoscale clustering of FcεRI receptors after Ag crosslinking where ordered regions are locally stabilized around the receptors making the distal regions more disordered. Such stabilization leads to an increase and decrease of diffusion coefficient of EGFP-GG, and PM-EGFP, respectively. These contrasting effects of LEX and FcεRI nanoclustering led us to hypothesize that the order in the outer leaflet changes globally (i.e., on the entire leaflet as opposed to local nanoscopic regions) due to LEX.

To test this hypothesis, we first monitored the diffusion of FAST-DiO, a disorder-preferring lipid probe (3), in the outer leaflet under various Lipid_ex_-LEX conditions. In control cells, the *D*_*av*_ of FAST-DiO is 0.79 ± 0.006 μm^2^/s (**Table 1**), close to the literature report of similar probes (55). We monitored the changes of the outer leaflet diffusion after DPPC- and DSPC-LEX. The *D*_*av*_ of FAST-DiO, similar to the inner leaflet probes, drops significantly under these conditions (Table 1) which is also reflected in the left-shift of the *D* CDFs (Fig. 3A). The *D*_Lex_/*D*_Ctrl_ values of FAST-DiO for DPPC- and DSPC-LEX are 0.42 and 0.28, respectively (Fig. 3B and Table 1). It appears that incorporation of order promoting lipids (DPPC or DSPC) makes the outer leaflet significantly more ordered.

**Figure 3.**
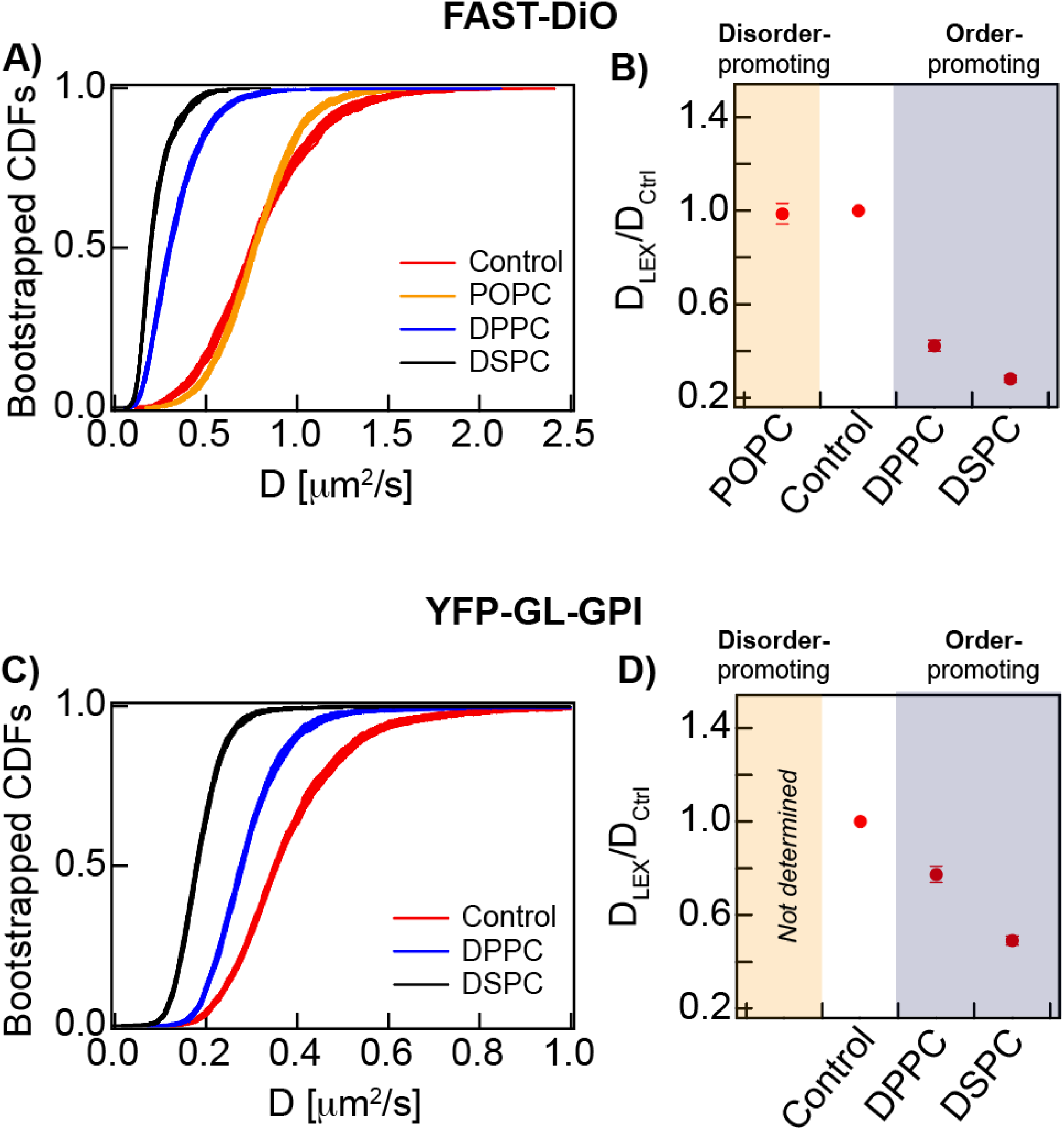
LEX-induced changes of outer leaflet order changes diffusion coefficient correspondingly in this leaflet. A) Bootstrapped cumulative distribution function (CDF) of diffusion coefficients (*D*) of FAST-DiO in control and LEX cells. B) The ratio of average *D* in LEX cells to that in control cells (*D*_Lex_/*D*_Ctrl_) for FAST-DiO for the indicated LEX. C-D) Bootstrapped CDFs of *D* and *D*_Lex_/*D*_Ctrl_ of YFP-GL-GPI on LEX and control cells. The error bars in the *D*_Lex_/*D*_Ctrl_ plots show the entire range of values of this parameter.

This is further supported from the modulation of diffusion behavior of yellow fluorescent protein (YFP)-tagged glycosylphosphatidylinositol (GPI) lipid anchor (YFP-GL-GPI (56)), an order-preferring outer leaflet lipid probe, in response to various LEX. The *D*_*av*_ of YFP-GL-GPI in the control cells is 0.38 ± 0.002 μm^2^/s (Table 1) which is similar to the previously reported values in RBL cell line and other cell lines (11, 12, 55, 57). *D* values for YFP-GL-GPI are slower in both DPPC- and DSPC-LEX cells (Fig. 3C). The *D*_*av*_ for DPPC- and DSPC-LEX cells are 0.29 ± 0.001 μm^2^/s and 0.19 ± 0.001 μm^2^/s, respectively, yielding corresponding *D*_Lex_/*D*_Ctrl_ values of 0.77 and 0.49 (Fig. 3D and Table 1). Interestingly, when we compare the *D*_Lex_/*D*_Ctrl_ values from the same Lipid_ex_, the effects of order-promoting lipids is more prominent on the diffusion of FAST-DiO (disorder-preferring probe) than YFP-GL-GPI (order-preferring probe) (Table 1). This might imply that the order-promoting Lipid_ex_ (DPPC and DSPC) have stronger impact in disordered regions than in ordered regions.

We confirmed that high concentrations of saturated lipids do not induce large gel-like domains in the outer leaflet, which, if formed, would be expected to result in large immobile fractions of order-preferring probes. We conducted fluorescence recovery after photobleaching (FRAP) on DSPC-LEX cells expressing YFP-GL-GPI (Fig. S4). While we observed slower florescence recovery (slower diffusion) than the control cells, we did not see any change in the immobile fraction. These results are consistent with the absence of large gel-like domains in DSPC-LEX cells. Together, our results on the diffusion behavior of various outer leaflet probes show that LEX induced changes of order in this leaflet correspond to whether Lipid_ex_ promotes membrane order or disorder.

It was more difficult to define the effects of POPC-LEX on the outer leaflet, and we could not use YFP-GL-GPI as a probe after LEX with POPC. TIRFM imaging showed non-uniform fluorescence distribution of YFP-GL-GPI in POPC-LEX cells (Fig S5, left panel). Quasi-TIRFM images further revealed that POPC-LEX depletes YFP-GL-GPI from the plasma membrane, relocating to internal membranes. By contrast, no obvious depletion of plasma membrane expression or internal membrane expression of YFP-GL-GPI was observed after LEX with order-promoting lipids (DPPC or DSPC). Both Quasi-TIRFM and TIRFM images of the inner leaflet probe, PM-EGFP, in POPC-LEX conditions does not show any obvious internalization of this probe (Fig S5, right panel). This implies that POPC-LEX induced internalization is likely to be restricted to only some outer leaflet components (see Discussion).

Measurements on outer leaflet diffusion after POPC-LEX could be made using FAST-DiO. As shown in Fig. 3A, the *D*_*av*_ of this probe remains unchanged after POPC-LEX resulting in a *D*_Lex_/*D*_Ctrl_ value of 0.99 (Fig. 3B and Table 1). This suggests that the physical properties of outer leaflet regions explored by FAST-DiO remain unchanged after POPC-LEX. However, when we measured the diffusion of alexa fluor 488 labelled cholera toxin B (AF488-CTxB) bound to GM1, another well-established outer leaflet marker that is order-preferring (50), we found the *D*_*av*_ of AF488-CTxB-GM1 (0.15 ± 0.001 μm^2^/s in control cells) increases to 0.24 ± 0.001 μm^2^/s after POPC-LEX (*D*_Lex_/*D*_Ctrl_ = 1.60; Fig. S3 and Table 1). Thus, POPC-LEX appears to make ordered regions less ordered or perhaps to make the outer leaflet overall more disordered as sensed by AF488-CTxB-GM1 whereas the properties of the disordered region remain unchanged as sensed by FAST-DiO.

### Stimulated Syk recruitment is enhanced when more ordered plasma membrane is created in the resting state

Recruitment of Syk kinase to the plasma membrane is the first step after FcεRI phosphorylation by Lyn kinase in mast cell signaling. We evaluated this process by measuring recruitment kinetics and density distribution of YFP-Syk using quantitative TIRF imaging. We sensitized the cells with unlabeled IgE and carried out LEX, which diffusion measurements show modifies inner leaflet order. Cells were stimulated by Ag-clustering of IgE-FcεRI.

We first imaged IgE-sensitized RBL cells transiently expressing YFP-Syk using TIRFM, which has an evanescent excitation wave that illuminates the surface attached plasma membrane and a small membrane-proximal cytosolic plane. In this resting condition, we observed a featureless diffuse distribution of YFP-Syk in the cytoplasm (58, 59). We then directly added Ag to the cells and imaged a representative cell in an imaging dish for each Lipid_ex_-LEX sample tested at 1 frame/min speed over 15 min after stimulation. The recruitment of YFP-Syk is clearly observed after 4-5 min as distinct punctate features against the cytosolic background (Supplemental Movie 1). These membrane-localized features appearing after stimulation are similar to those reported previously (59). We also determined the density of YFP-Syk puncta (i.e., number of puncta/cell area) as function of time. We found that the puncta density reaches a steady-state by 15 min after stimulation (Fig. 4A), and we proceeded to image many cells at this stimulated steady-state to obtain a distribution of the density of plasma membrane-localized YFP-Syk puncta (Fig. 4B).

**Figure 4.**
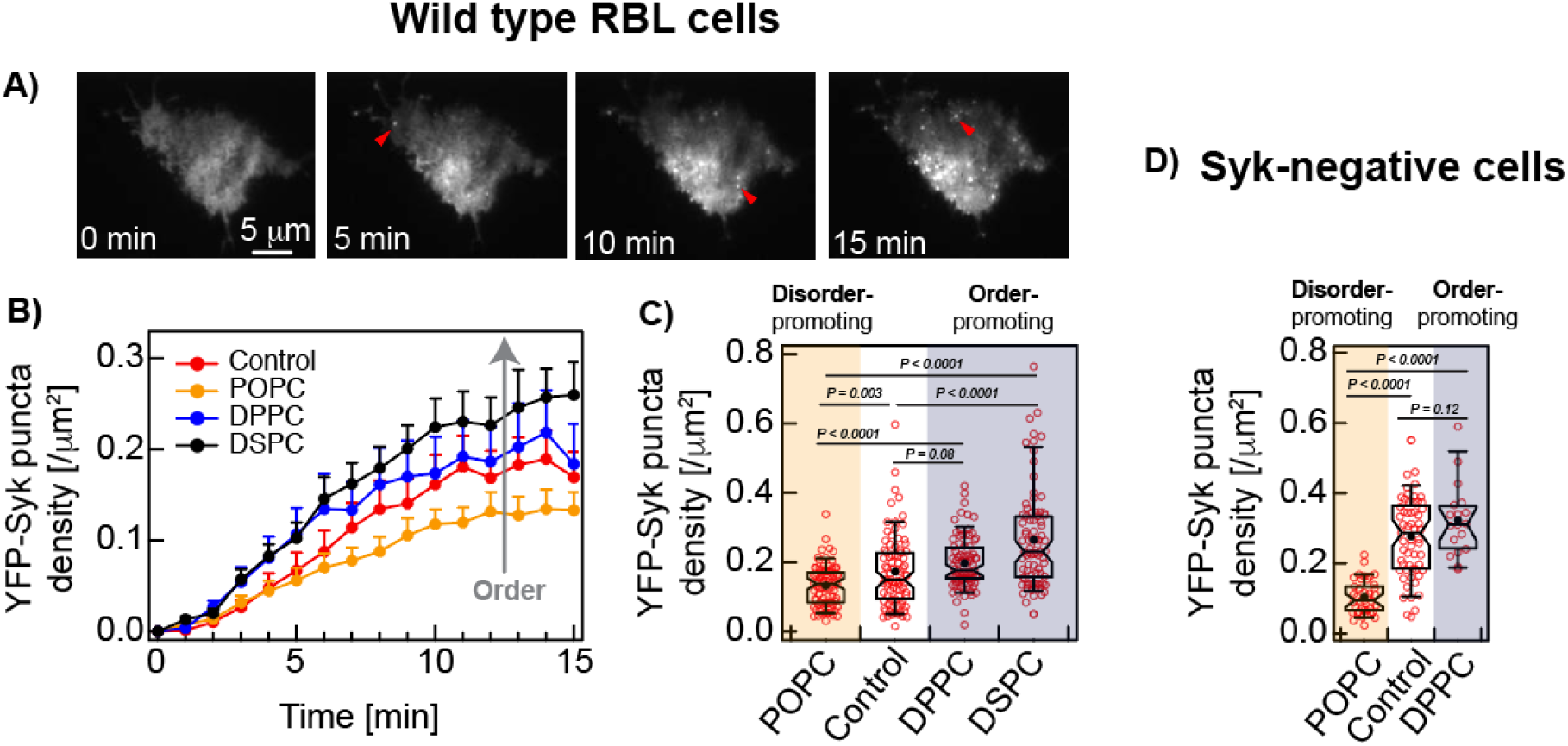
Stimulated recruitment of YFP-Syk to the inner leaflet of the plasma membrane correlates with the resting state order of this leaflet modulated by outer leaflet LEX. A-B) Kinetics of YFP-Syk recruitment at different exchanged lipids and control conditions in wildtype RBL cells. Snapshots of TIRF a representative control cell expressing YFP-Syk at various time points after the addition of antigen (Ag), DNP-BSA, shown in A. Number of cells = 12-18 for each data point. The error bars represent standard error of the mean (SEM). B) Box plots of density of the recruited YFP-Syk punctas in stimulated steady-state (15-30 min after stimulation). Number of cells > 80 for each condition. C) Box plots of density of YFP-Syk punctas in Syk-negative cells (transiently expressing YFP-Syk) after 15 min stimulation. Number of cells >15 at each condition. Box height corresponds to 25th to 75th percentile; error bars represent 9th to 91st percentile of entire dataset; mean and median values are represented as solid circle and bar, respectively; notches signify 95% CI of the median.

Fig. 4B shows the time course of stimulated YFP-Syk recruitment for control cells compared to those with more ordered (DPPC-LEX or DSPC-LEX) or disordered (POPC-LEX) outer leaflet compositions (and correspondingly more ordered or disordered inner leaflet due to inter-leaflet coupling) as established under resting conditions (Figs. 2 and 3). In the absence of Ag-stimulation, we observed negligible YFP-Syk recruitment in control cells and with Lipid_ex_ tested. For stimulated cells, the rate of increase of YFP-Syk puncta density is similar for control and LEX cells. However, the YFP-Syk puncta density at the stimulated steady-state (15 min post-stimulation) is remarkably different across the Lipid_ex_ tested. YFP-Syk puncta density is significantly lower for POPC-LEX cells compared to control cells. We observed the reverse for cells exchanged with order-promoting lipids (DPPC-LEX and DSPC-LEX). Following the same trend as the inter-leaflet effects on diffusion (Figs. 2B and D), we further observed that DSPC-LEX has greater impact than DPPC-LEX on stimulated YFP-Syk recruitment.

To gain more quantitative and statistically robust information, we imaged ∼100 cells expressing YFP-Syk in stimulated steady-state for control cell and each Lipid_ex_ tested. We found that ∼80% of all cells show stimulated YFP-Syk puncta in all cases. However, the puncta density of LEX cells is statistically different from the control cells (Fig. 4B). Compared to control cells (median YFP-Syk density = 0.15 /μm^2^), cells exchanged with order-promoting lipids, DPPC-LEX and DSPC-LEX show 18% and 54% higher median YFP-Syk density, respectively, at stimulated steady-state, whereas cells exchanged with disorder-promoting lipid, POPC-LEX, shows 8% lower median YFP-Syk density. Overall, the degree of stimulated Syk recruitment, a plasma-membrane proximal signaling event, appears to correlate with the inner leaflet membrane order created in resting cells.

To consider the possibility that Syk endogenously expressed in RBL cells affects stimulated recruitment of transiently expressed YFP-Syk, we employed Syk-negative cells (37). This RBL variant cell line expresses no detectable Syk and correspondingly has an abrogated signaling cascade when stimulated by Ag. We transiently transfected YFP-Syk into Syk-negative cells and tested effects of POPC-LEX (disorder-promoting) and DPPC-LEX (order-promoting) compared with control cells. We found for control cells that the stimulated steady-state density of plasma membrane localized YFP-Syk puncta is generally larger in the Syk-negative cells compared to the wild type RBL cells (median values are 0.29 and 0.15 /μm^2^, respectively), as might be expected from eliminated competition with endogenous Syk. In Syk-negative cells, the steady-state levels of recruited YFP-Syk puncta are significantly lower for POPC-LEX compared to control cells, and DPPC-LEX are significantly higher than POPC-LEX (Fig. 4C). We were unable to evaluate DSPC-LEX because the Syk-negative cells did not sufficiently survive this treatment. Overall, the trends of stimulated YFP-Syk recruitment are qualitatively similar for wild type and Syk-negative RBL cells (Figs. 4B and C): LEX to increase disorder-promoting lipids in the outer leaflet decreases stimulated Syk recruitment, while this signaling activity is enhanced by LEX to increase order-promoting lipids in the outer leaflet. These results are consistent with inter-leaflet coupling and participation of inner leaflet order in the process leading to Syk recruitment.

### More ordered resting plasma membrane results in stronger stimulated degranulation

Stimulated Syk recruitment is a signaling event proximal to antigen-clustering of IgE-FcεRI in the membrane. We also tested whether modulation of outer leaflet order correlates with the culminating step of the signaling cascade, which is exocytosis of secretory granules containing β-hexosaminidase and other chemical mediators. As shown in Fig. 5, we observed a similar trend for antigen-stimulated degranulation: POPC-LEX to increase disorder-promoting lipids in the outer leaflet decreases stimulated degranulation by almost 80% compared to control cells, whereas DPPC-LEX and DSPC-LEX to increase order-promoting lipids in the outer leaflet increase this response by 30% and 50%, respectively. Therefore, stimulated degranulation correlates with LEX-altered resting state membrane order in a fashion analogous to stimulated Syk recruitment.

**Figure 5.**
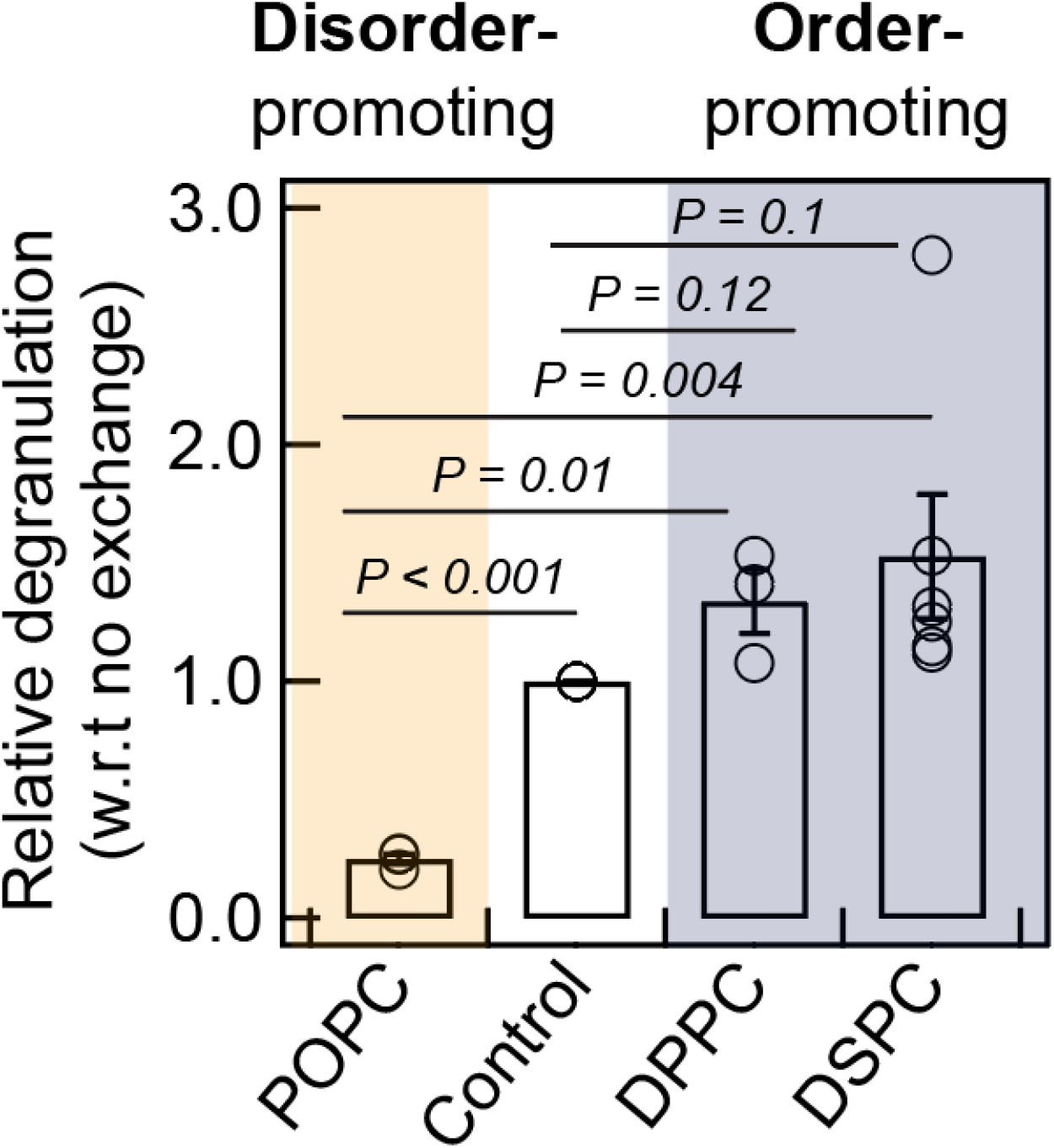
Degree of stimulated degranulation correlates with resting state plasma membrane order modulated by outer leaflet LEX. Relative degranulation for specified Lipidex-LEX conditions are shown. The error bars are standard error of mean (SEM).

## DISCUSSION

### LEX is a useful approach for studying inter-leaflet coupling in live cells

Inter-leaflet coupling between compositionally distinct inner and outer leaflets is a key biophysical determinant of steady-state plasma membrane heterogeneity (30, 60, 61). Transmembrane (TM) protein-based coupling may occur when the local membrane environment of these proteins is longitudinally coupled. For example, some palmitoylated TM proteins locally associate with ordered domains (62). Lipid-based coupling is reported to be governed by (transient) immobilization of long-chain lipids either by extracellular cross-linking (for example, by multivalent toxins (21)) or by intracellular coupling by cortical actin machinery (20)). These experimental strategies to demonstrate lipid-based inter-leaflet coupling chiefly depend on crosslinking or immobilizing a component in one leaflet and monitoring the effect on the opposing leaflet.

Here we focus on an alternative mechanism whereby we exploit LEX to manipulate lipid composition of outer leaflet, and we measure the diffusion properties of inner leaflet by highly sensitive ImFCS measurements of diffusion. We further evaluate mast cell signaling through FcεRI to test functional consequences of LEX. The exchange of outer leaflet lipids with exogeneous lipid (Lipid_ex_) by LEX offers a straightforward method to incorporate selected lipids. The mαCD-Lipid_ex_ complex acts as a lipid donor while uncomplexed mαCD acts as a lipid acceptor that removes endogenous outer leaflet lipids. Such donor-acceptor lipid exchange systems using various types of cyclodextrin (CD) have been widely used to create asymmetric artificial lipid membranes (60). This provides an excellent experimental platform to study inter-leaflet coupling in these model systems. Applications of mαCD has some advantages over other CD species for experiments using living cells. The cavity size of mαCD is not appropriate to make complex with cholesterol. Therefore, mαCD-catalyzed lipid exchange does not significantly change cellular cholesterol content, unlike methyl-β-cyclodextrin (mβCD) catalyzed lipid exchange protocols which extracts cholesterol from the plasma membrane (63). Another favorable property is that mαCD has low specificity for the lipid structures allowing efficient extraction of various non-sterol lipids (e.g., PC and SM) from the outer leaflet.

The changes of outer leaflet biophysical properties due to LEX shows interesting trends. The *D*_*av*_ of FAST-DiO, a disordered preferring probe, does not change after POPC-LEX (Fig. 3A). If the outer leaflet has pre-existing heterogeneity with both more disordered and less disordered regions, we reasoned that POPC, a disorder-promoting lipid, replaces endogenous lipids in the disordered regions of the outer leaflet into which FAST-DiO largely partitions, and this *like-for-like* replacement in the membrane environment has little impact on the diffusion in this region. In contrast, POPC_ex_ replacement of order-promoting endogenous lipids likely decreases membrane order in more ordered regions of the plasma membrane outer leaflet. This appears to be reflected in the diffusion change of AF488-CTxB-GM1 (Fig. S3 and Table 1). On the other hand, because CTxB can induce ordered domain formation (23, 64), another interpretation is that replacement of sphingolipids and other order-promoting lipids with POPC prevents formation of such domains. It should be noted the slower diffusion of CTxB relative to FAST-DiO in the absence of LEX could reflect the complexation of CTxB with up to five copies of GM1 as well as localization in domains of increased membrane order. The decrease of *D*_*av*_ for both disorder- and order-preferring probes (FAST-DiO and YFP-Gl-GPI) after DPPC- and DSPC-LEX (Figs. 3A and 3C respectively) indicates changes in membrane order throughout outer leaflet (global change). The observed stronger changes in the diffusion of FAST-DiO than that of YFP-GL-GPI (i.e., *D*_Lex_/*D*_Ctrl_ values were <0.3 for FAST-DiO and >0.3 for YFP-GL-GPI; Table 1) further suggest that the disordered regions of the outer leaflet are more strongly influenced by the incorporation of order-promoting lipids.

As for the robustness of LEX, it has been successfully applied on multiple cell lines including RBC, CHO, HeLa, RBL, macrophages, and A549 (31, 32, 40, 65). Although here we only used phospholipids as exogenous lipids, sphingolipids can similarly be incorporated (32). The effect of LEX on particular plasma membrane-localized processes also can be tested. Examples include stimulated phosphorylation of insulin receptor in CHO cells (40), endocytosis of model nanoparticles in A549 cells (66), and stimulated Syk recruitment during mast cell signaling (this study). This demonstrates the robustness and usefulness of this strategy. It however should be noted that degree of incorporation can depend on both the types of cells and exogenous lipid species (33). Therefore, the degree of lipid exchange should be quantified, for example by TLC as in Fig. S1 or mass spectrometry (66), as part of interpreting the biophysical and functional results.

Live cells, not surprisingly, respond to compensate for changes accompanying such substantial amount (50-100%) of exogenous lipid exchange. For example, the diffusion of AF488-IgE-FcεRI is restored over two hours after LEX (Fig. S2). Cells may either degrade or remodel the exogenous lipids (31, 32, 67), and replenish exchanged lipids by synthesizing new lipids or delivering pre-existing lipids to the plasma membrane from internal sources. Therefore, experiments must be restricted to times in which LEX-induced composition is stable, which can depend on the choice of lipid (32). Some unintended indirect effects of LEX should be considered. For example, YFP-GL-GPI is depleted from the plasma membrane in POPC-LEX cells (Fig. S5). This was not observed for DPPC- or DSPC-LEX. Although investigation of the mechanisms underlying YFP-GL-GPI depletion is beyond the scope of this work, one possibility is LEX might trigger internalization of the outer leaflet components of ordered regions such as YFP-GL-GPI. For example, a high fraction of saturated lipids, a key component of ordered regions, is exchanged with POPC rendering the ordered regions in the outer leaflet sparse, unstable or both. This might affect constitutive endocytosis and recycling, both of which would halt when a step of membrane trafficking is blocked. Rafaei et al. showed that the ordered regions of the plasma membrane strongly affect the recycling process in the resting CHO cells (68). In addition, replacing order-stabilizing sterols with sterols that do not support ordered nanodomains slows membrane trafficking as judged by inhibition of endocytosis (34). In the current context, the loss of ordered regions in the POPC-exchanged plasma membrane may disrupt YFP-GL-GPI recycling causing them to concentrate in the internal membranes or lysosomes for degradation. We cannot rule out other indirect effects of LEX, including changes in cholesterol distribution within the plasma membrane or its interactions with the actin cytoskeleton.

### LEX induces global changes in the inner leaflet diffusion properties

Our recent high resolution studies (12, 14) showed that crosslinking of IgE-FcεRI by multivalent Ag stabilizes phase-like separation in the inner leaflet. Proximal to the clustered receptors is an ordered environment, which is surrounded by a disordered region. In other words, the ordered and disordered regions become more distinctive in the stimulated steady-state, and this is reflected in the diffusion behavior of order- and disorder-preferring lipid probes of the inner leaflet, PM-EGFP and EGFP-GG, respectively. Consistent with the probes spending more time in their preferred membrane regions, the diffusion coefficient (*D*) of PM-EGFP decreases while that of EGFP-GG decreases in Ag-stimulated relative to resting cells (12). We observed the same trend in the diffusion behavior of these probes in resting cells after inhibition of actin polymerization by cytochalasin D (25), which is consistent with cytoskeletal regulation of phase-like properties (69).

In contrast, the diffusion coefficient of both PM-EGFP and EGFP-GG increases when the outer leaflet is exchanged with disorder-promoting lipid, POPC, and their diffusion coefficients decrease when outer leaflet lipids are exchanged with order-promoting lipids, DPPC or DSPC (Fig. 2). Interestingly, the *D*_Lex_/*D*_Ctrl_ values of both PM-EGFP and EGFP-GG are similar for each of these LEX conditions (Figs. 2B and D). This suggests that any preexisting ordered and disordered regions in the inner leaflet do not become more distinctive due to LEX driven inter-leaflet interactions. Instead, both regions appear to become almost equally more disordered when a disorder-promoting lipid (POPC) is introduced in the outer leaflet. When order-promoting lipids (DPPC or DSPC) is introduced in the outer leaflet, the inner leaflet appears to become globally more ordered as reflected by increased diffusion coefficients of both inner leaflet probes. Notably, DSPC lipid vesicles having higher transition temperature than DPPC lipid vesicles (54°C and 41°C respectively) is likely to induce a stronger ordering effect than DPPC and correspondingly, *D*_Lex_/*D*_Ctrl_ of all probes tested here is smaller in DSPC-LEX cells than in DPPC-LEX cells.

An alternate explanation for the changes in diffusion in the inner leaflet is that they are due to a change in inner leaflet lipid composition induced by LEX rather than due to interleaflet coupling. Several lines of evidence indicate this is unlikely. First, in both artificial lipid vesicles and cells, LEX is restricted to outer leaflet lipids (31, 70, 71). Second, the lipids used for LEX undergo only very slow spontaneous lipid flip-flop (xyr-73). Third, after LEX the lipid introduced into the outer leaflet remains largely exposed to removal by a second round of lipid exchange for 1 hour, indicating that they remain in the outer leaflet (31, 32). Nevertheless, we cannot rule out the possibility that there is some small change in inner leaflet lipid composition due directly to LEX.

Although we did not measure an order parameter of the inner leaflet directly, the diffusion of these lipid probes is likely to be inversely related to membrane order (56, 57). Both PM-EGFP and EGFP-GG showed fastest diffusion in POPC-LEX cells and slowest diffusion in DSPC-LEX cells among the conditions tested here (Fig. 2 and Table 1). This implies that the inner leaflet is most disordered in POPC-LEX cells and most ordered in DSPC-LEX cells. For the Lipid_ex_ we tested, the apparent inner leaflet order correlates with order-preference: POPC < control < DPPC < DSPC.

The observation of global changes of diffusion of both order- and disorder-preferring lipid probes after LEX resembles those we observed previously after antibody crosslinking and immobilization of CTxB-GM1 complexes (74). However, the degree of probe diffusion changes in these two treatments (LEX and GM1 crosslinking) are different. The *D* changes from PM-EGFP and EGFP-GG were ∼32% and 55% after DPPC- and DSPC-LEX (Figs. 3D and 3H). In contrast, antibody crosslinking of GM1 with anti-CTxB antibody results in only ∼16% reduction of *D* of these probes (74). LEX appears to cause changes in the order of the entire outer leaflet. In contrast, crosslinking GM1 directly affects a single membrane component. Thus, this difference is likely due to relatively larger inter-leaflet coupling effects (as monitored by inner leaflet probe diffusion) by LEX than antibody crosslinking of GM1. Overall, our group so far has experimentally evaluated various modes of ‘outside-in’ inter-leaflet interactions (this study and (12, 74)) and found primarily two different outcomes as measured from precise diffusion measurements of these inner leaflet probes: either stabilization of phase-like separation (e.g., Ag-clustering of IgE-FcεRI receptors (12) or global change of membrane order (LEX in this study) and antibody crosslinking of CTxB-GM1 (74)). This body of work establishes the power of our bootstrapping-based analysis of diffusion coefficients of lipid probes measured by ImFCS (12) to delineate subtle changes of membrane organization in live cells and thereby reveal underlying biophysical principles.

LEX mediated inter-leaflet coupling demonstrated here is fundamentally distinctive from other proposed ‘outside-in’ mechanisms which generally involve immobilization of membrane components. It was reported that immobilization of outer leaflet GM1 or GPI-anchored protein by antibody crosslinking leads to the changes in the diffusion properties of inner leaflet probes (21, 75). Likewise, the Mayor group recently showed that crosslinking and immobilization of saturated, long-chain ceramides, but not their unsaturated counterpart, induces ordered regions in the inner leaflet (23). In contrast, LEX does not immobilize any membrane component. We therefore infer that lipid-mediated inter-leaflet interactions do not strictly require immobilization of a membrane component. The key effect of crosslinking may be local formation of a more ordered state rather than the immobilization that also accompanies crosslinking.

### More ordered membrane in the poised resting state leads to stronger functional response

Ag-stimulated stabilization of ordered regions has been demonstrated to play a key role in facilitating receptor phosphorylation above non-functional threshold in IgE-FcεRI mediated cell signaling (5, 12, 14, 53, 76, 77). Preferential partitioning of Lyn kinase with Ag-clustered receptors in the ordered regions facilitates net receptor phosphorylation and suppresses dephosphorylation TM phosphatases, which prefer the disordered regions. Phosphorylated FcεRI then recruit Syk kinase to propagate intracellular signaling cascade leading to cellular degranulation.

We monitored, for different Lipid_ex_, stimulated recruitment of YFP-Syk kinase to the inner leaflet of the plasma membranes. Our time-lapse TIRFM imaging results clearly demonstrate that the density of plasma membrane localized YFP-Syk in Ag-stimulated steady-state correlates with the order-preference of Lipid_ex_ and membrane order detected by inner leaflet diffusion probes. We observed the same trend for Ag-stimulated degranulation. Notably, we did not detect Syk recruitment in the absence of Ag-stimulation even if the inner leaflet is more ordered after LEX with order-promoting lipids. This means that increasing membrane order does not by itself cause sufficiently strong colocalization of Lyn kinase and FcεRI phosphorylation. Our results therefore suggest that a more ordered resting membrane enhances stimulation-induced stabilization of phase-like separation which is required for FcεRI phosphorylation and consequent Syk recruitment. Simply stated, ordered and disordered regions in the inner leaflet appear to be more distinctive in the stimulated steady-state when this leaflet is more ordered in the resting conditions. More distinctive regions would provide better spatial protection to the crosslinked FcεRI receptors from dephosphorylation by disorder-preferring TM phosphatases. The impact of altered Syk recruitment (an early signaling event) driven by changes in membrane order seems to propagate to the degree of degranulation (ultimate cellular response) (Fig. 5) although there are many signaling steps in between, and membrane fusion involved in exocytosis may also be affected by membrane order.

Alternative scenarios leading to stronger or weaker Syk recruitment depending on resting state inner leaflet order may also be possible. For example, Lyn’s kinase activity may be altered depending on the surrounding membrane environment. Young et al. showed that specific activity of Lyn isolated from ordered regions is about five times higher than that isolated from the disordered region (76). Along that line, it was recently shown that incorporation of highly unsaturated lipid, ω-3 fatty acid, in the plasma membrane inhibits mast cell functions *in vivo* (78), and the authors suggest that this is related to inhibition of Lyn kinase activity. The high amount of ω-3 fatty acid, similar to that of POPC incorporation by LEX, is likely to make membrane more disordered which may reduce the kinase activity of Lyn. Also, the conformation of Ag-clustered receptors may be altered in POPC-LEX conditions leading to cytosolic Tyr residues becoming less accessible for phosphorylation and subsequent binding to Syk kinase, and/or more accessible to disorder-preferring TM phosphatase. Such conformational changes depending on the lipid environment was recently proposed for ligand-stimulated insulin receptors (39). Finally, we note that correlating two stimulated functional read-outs (Syk recruitment and degranulation) with membrane order does not speak to the influence of membrane order on other intermediate signaling processes (for example, stimulated calcium mobilization).

## CONCLUSIONS

This study quantitatively evaluates inter-leaflet coupling in live cells by measuring diffusion changes of inner leaflet lipid probes resulting from the perturbation of outer leaflet lipid composition by LEX. Our results provide compelling evidence of a novel, lipid-based inter-leaflet coupling mechanism that does not necessarily involve long-chain (>20 carbon) lipids or immobilization of a membrane component. The effect appears to be global: increasing order-promoting lipids in the outer leaflet increases the order of the inner leaflet, and increasing the content of disorder-preferring lipids in the outer leaflet decreases the order of the inner leaflet. There are clear functional consequences of inter-leaflet coupling induced by LEX as demonstrated with Ag-stimulated, IgE-FcεRI mediated mast cell signaling where membrane order is known to regulate protein interactions associated with TM signaling events. Overall, we developed an experimental platform combining LEX, ImFCS, and quantitative fluorescence imaging to evaluate functional, lipid-based inter-leaflet coupling of plasma membranes in live cells.

## AUTHOR CONTRIBUTIONS

BAB, EL, and NB conceived the project, G-SY, AWW, BY, PS, and NB performed the experiments, all authors analysed data, BAB, EL, and NB wrote the manuscript drafts with all authors critically contributing to the final manuscript.

## ACKNOWLEDGEMENTS

We thank Dr. Alex Batrouni and Prof. Jeremy Baskin (Cornell University) for assisting with the FRAP measurements, and Dr. David Holowka for helpful discussion. This work is supported by National Institute of General Medical Sciences (NIGMS) Grant R01GM117552 to B.A.B. and R35GM122493 to E.L. The content is solely the responsibility of the authors and does not necessarily represent the official views of NIGMS or NIH.

## REFERENCES

1. M. Dykstra, A. Cherukuri, H. W. Sohn, S. J. Tzeng, S. K. Pierce, Location is everything: lipid rafts and immune cell signaling. Annu Rev Immunol 21, 457–481 (2003).

2. T. Baumgart et al., Large-scale fluid/fluid phase separation of proteins and lipids in giant plasma membrane vesicles. Proc Natl Acad Sci U S A 104, 3165–3170 (2007).

3. E. Sezgin et al., Elucidating membrane structure and protein behavior using giant plasma membrane vesicles. Nat Protoc 7, 1042–1051 (2012).

4. P. Sengupta, A. Hammond, D. Holowka, B. Baird, Structural determinants for partitioning of lipids and proteins between coexisting fluid phases in giant plasma membrane vesicles. Biochim Biophys Acta 1778, 20–32 (2008).

5. E. Sezgin, I. Levental, S. Mayor, C. Eggeling, tThe mystery of membrane organization: composition, regulation and roles of lipid rafts. Nat Rev Mol Cell Biol 18, 361–374 (2017).

6. F. M. Goni, “Rafts”: A nickname for putative transient nanodomains. Chem Phys Lipids 218, 34–39 (2019).

7. C. Eggeling et al., Direct observation of the nanoscale dynamics of membrane lipids in a living cell. Nature 457, 1159–1162 (2009).

8. A. Honigmann et al., Scanning STED-FCS reveals spatiotemporal heterogeneity of lipid interaction in the plasma membrane of living cells. Nat Commun 5, 5412 (2014).

9. D. M. Owen, D. J. Williamson, A. Magenau, K. Gaus, Sub-resolution lipid domains exist in the plasma membrane and regulate protein diffusion and distribution. Nat Commun 3, 1256 (2012).

10. P. R. Nicovich, J. M. Kwiatek, Y. Ma, A. Benda, K. Gaus, FSCS Reveals the Complexity of Lipid Domain Dynamics in the Plasma Membrane of Live Cells. Biophysical Journal 114, 2855–2864 (2018).

11. N. Bag, D. A. Holowka, B. A. Baird, Imaging FCS delineates subtle heterogeneity in plasma membranes of resting mast cells. Mol Biol Cell 31, 709–723 (2020).

12. N. Bag et al., Lipid-based and protein-based interactions synergize transmembrane signaling stimulated by antigen clustering of IgE receptors. Proc Natl Acad Sci U S A 118, e2026583118 (2021).

13. S. A. Shelby, D. Holowka, B. Baird, S. L. Veatch, Distinct stages of stimulated FcepsilonRI receptor clustering and immobilization are identified through superresolution imaging. Biophys J 105, 2343–2354 (2013).

14. S. A. Shelby, S. L. Veatch, D. A. Holowka, B. A. Baird, Functional nanoscale coupling of Lyn kinase with IgE-FcepsilonRI is restricted by the actin cytoskeleton in early antigen-stimulated signaling. Mol Biol Cell 27, 3645–3658 (2016).

15. T. Kambayashi, G. A. Koretzky, Proximal signaling events in Fc epsilon RI-mediated mast cell activation. J Allergy Clin Immunol 119, 544-552; quiz 553-544 (2007).

16. R. M. Young, X. Zheng, D. Holowka, B. Baird, Reconstitution of regulated phosphorylation of FcepsilonRI by a lipid raft-excluded protein-tyrosine phosphatase. J Biol Chem 280, 1230–1235 (2005).

17. J. Rivera, A. M. Gilfillan, Molecular regulation of mast cell activation. J Allergy Clin Immunol 117, 1214-1225; quiz 1226 (2006).

18. J. H. Lorent et al., Plasma membranes are asymmetric in lipid unsaturation, packing and protein shape. Nat Chem Biol 16, 644–652 (2020).

19. T.-Y. Wang, J. R. Silvius, Cholesterol Does Not Induce Segregation of Liquid-Ordered Domains in Bilayers Modeling the Inner Leaflet of the Plasma Membrane. Biophysical Journal 81, 2762–2773 (2001).

20. R. Raghupathy et al., Transbilayer lipid interactions mediate nanoclustering of lipid-anchored proteins. Cell 161, 581–594 (2015).

21. I. Koyama-Honda et al., High-speed single-molecule imaging reveals signal transduction by induced transbilayer raft phases. Journal of Cell Biology 219, e202006125 (2020).

22. S. Chiantia, E. London, Acyl chain length and saturation modulate interleaflet coupling in asymmetric bilayers: effects on dynamics and structural order. Biophys J 103, 2311–2319 (2012).

23. S. Arumugam et al., Ceramide structure dictates glycosphingolipid nanodomain assembly and function. Nat Commun 12, 3675 (2021).

24. G. van Meer, Dynamic transbilayer lipid asymmetry. Cold Spring Harb Perspect Biol 3, a004671 (2011).

25. T. Kobayashi, A. K. Menon, Transbilayer lipid asymmetry. Current Biology 28, PR386–R391 (2018).

26. Q. Lin, E. London, Ordered raft domains induced by outer leaflet sphingomyelin in cholesterol-rich asymmetric vesicles. Biophys J 108, 2212–2222 (2015).

27. T. A. Enoki, G. W. Feigenson, Improving our picture of the plasma membrane: Rafts induce ordered domains in a simplified model cytoplasmic leaflet. Biochim Biophys Acta Biomembr 1864, 183995 (2022).

28. J. W. St Clair, S. Kakuda, E. London, Induction of Ordered Lipid Raft Domain Formation by Loss of Lipid Asymmetry. Biophys J 119, 483–492 (2020).

29. S. Kakuda, P. Suresh, G. Li, E. London, Loss of plasma membrane lipid asymmetry can induce ordered domain (raft) formation. J Lipid Res 63, 100155 (2022).

30. M. Doktorova, J. L. Symons, I. Levental, Structural and functional consequences of reversible lipid asymmetry in living membranes. Nat Chem Biol 16, 1321–1330 (2020).

31. G. Li et al., Efficient replacement of plasma membrane outer leaflet phospholipids and sphingolipids in cells with exogenous lipids. Proc Natl Acad Sci U S A 113, 14025–14030 (2016).

32. G. Li et al., Replacing plasma membrane outer leaflet lipids with exogenous lipid without damaging membrane integrity. PLoS One 14, e0223572 (2019).

33. P. Suresh, E. London, Using cyclodextrin-induced lipid substitution to study membrane lipid and ordered membrane domain (raft) function in cells. Biochim Biophys Acta Biomembr 1864, 183774 (2021).

34. J. H. Kim, A. Singh, M. Del Poeta, D. A. Brown, E. London, The effect of sterol structure upon clathrin-mediated and clathrin-independent endocytosis. J Cell Sci 130, 2682–2695 (2017).

35. S. L. Veatch, S. L. Keller, A closer look at the canonical ‘raft mixture’ in model membrane studies. Biophysical Journal 84, 725–726 (2003).

36. A. J. Torres, L. Vasudevan, D. Holowka, B. A. Baird, Focal adhesion proteins connect IgE receptors to the cytoskeleton as revealed by micropatterned ligand arrays. Proceedings of the National Academy of Sciences of the United States of America 105, 17238–17244 (2008).

37. J. Zhang, E. H. Berenstein, R. L. Evans, R. P. Siraganian, Transfection of Syk protein tyrosine kinase reconstitutes high affinity IgE receptor-mediated degranulation in a Syk-negative variant of rat basophilic leukemia RBL-2H3 cells. J Exp Med 184, 71–79 (1996).

38. P. S. Pyenta, D. Holowka, B. Baird, Cross-correlation analysis of inner-leaflet-anchored green fluorescent protein co-redistributed with IgE receptors and outer leaflet lipid raft components. Biophys J 80, 2120–2132 (2001).

39. D. L. Wakefield, D. Holowka, B. Baird, The FcepsilonRI Signaling Cascade and Integrin Trafficking Converge at Patterned Ligand Surfaces. Mol Biol Cell 10.1091/mbc.E17-03-0208 (2017).

40. P. Suresh, W. T. Miller, E. London, Phospholipid exchange shows insulin receptor activity is supported by both the propensity to form wide bilayers and ordered raft domains. J Biol Chem 297, 101010 (2021).

41. R. M. Naal, J. Tabb, D. Holowka, B. Baird, In situ measurement of degranulation as a biosensor based on RBL-2H3 mast cells. Biosens Bioelectron 20, 791–796 (2004).

42. J. Schindelin et al., Fiji: an open-source platform for biological-image analysis. Nat Methods 9, 676–682 (2012).

43. N. Bag, J. Sankaran, A. Paul, R. S. Kraut, T. Wohland, Calibration and limits of camera-based fluorescence correlation spectroscopy: a supported lipid bilayer study. ChemPhysChem 13, 2784–2794 (2012).

44. J. Sankaran, N. Bag, R. S. Kraut, T. Wohland, Accuracy and precision in camera-based fluorescence correlation spectroscopy measurements. Anal Chem 85, 3948–3954 (2013).

45. J. Sankaran, X. Shi, L. Y. Ho, E. H. Stelzer, T. Wohland, ImFCS: a software for imaging FCS data analysis and visualization. Opt Express 18, 25468–25481 (2010).

46. E. K. Fridriksson et al., Quantitative analysis of phospholipids in functionally important membrane domains from RBL-2H3 mast cells using tandem high-resolution mass spectrometry. Biochemistry 38, 8056–8063 (1999).

47. A. Gupta, S. Muralidharan, F. Torta, M. R. Wenk, T. Wohland, Long acyl chain ceramides govern cholesterol and cytoskeleton dependence of membrane outer leaflet dynamics. Biochim Biophys Acta Biomembr 1862, 183153 (2020).

48. M. J. Swamy et al., Coexisting domains in the plasma membranes of live cells characterized by spin-label ESR spectroscopy. Biophys J 90, 4452–4465 (2006).

49. A. Gidwani, D. Holowka, B. Baird, Fluorescence anisotropy measurements of lipid order in plasma membranes and lipid rafts from RBL-2H3 mast cells. Biochemistry 40, 12422–12429 (2001).

50. S. A. Shelby, I. Castello-Serrano, K. C. Wisser, I. Levental, S. L. Veatch, Membrane phase separation drives organization at B cell receptor clusters. BioRxiv 10.1101/2021.05.12.443834 (2021).

51. D. Holowka, K. Thanapuasuwan, B. Baird, Short chain ceramides disrupt immunoreceptor signaling by inhibiting segregation of Lo from Ld Plasma membrane components. Biol Open 7 (2018).

52. K. A. Field, D. Holowka, B. Baird, FceRI-mediated recruitment of p53/56lyn to detergent-resistant membrane domains accompanies cellular signaling. Proc Natl Acad Sci U S A 92, 9201–9205 (1995).

53. K. A. Field, D. Holowka, B. Baird, Compartmentalized activation of the high affinity immunoglobulin E receptor within membrane domains. Journal of Biological Chemistry 272, 4276–4280 (1997).

54. A. M. Davey, R. P. Walvick, Y. Liu, A. A. Heikal, E. D. Sheets, Membrane order and molecular dynamics associated with IgE receptor cross-linking in mast cells. Biophys J 92, 343–355 (2007).

55. S. Huang, S. Y. Lim, A. Gupta, N. Bag, T. Wohland, Plasma membrane organization and dynamics is probe and cell line dependent. Biochim Biophys Acta 10.1016/j.bbamem.2016.12.009 (2016).

56. A. Pralle, P. Keller, E.-L. Florin, J. K. H. Hörbor, Sphingolipid–Cholesterol Rafts Diffuse as Small Entities in the Plasma Membrane of Mammalian Cells. J Cell Biol 148, 997–1007 (2000).

57. A. K. Kenworthy et al., Dynamics of putative raft-associated proteins at the cell surface. J Cell Biol 165, 735–746 (2004).

58. R. Das, S. Hammond, D. Holowka, B. Baird, Real-time cross-correlation image analysis of early events in IgE receptor signaling. Biophys J 94, 4996–5008 (2008).

59. S. L. Schwartz et al., Differential mast cell outcomes are sensitive to FcepsilonRI-Syk binding kinetics. Mol Biol Cell 28, 3397–3414 (2017).

60. E. London, Membrane Structure-Function Insights from Asymmetric Lipid Vesicles. Acc Chem Res 52, 2382–2391 (2019).

61. T. Fujimoto, I. Parmryd, Interleaflet Coupling, Pinning, and Leaflet Asymmetry-Major Players in Plasma Membrane Nanodomain Formation. Front Cell Dev Biol 4, 155 (2016).

62. I. Levental, K. R. Levental, F. A. Heberle, Lipid Rafts: Controversies Resolved, Mysteries Remain. Trends Cell Biol 30, 341–353 (2020).

63. R. Zidovetzki, I. Levitan, Use of cyclodextrins to manipulate plasma membrane cholesterol content: evidence, misconceptions and control strategies. Biochim Biophys Acta 1768, 1311–1324 (2007).

64. A. T. Hammond et al., Crosslinking a lipid raft component triggers liquidordered–liquid disordered phase separation in modelplasma membranes. Proc. Natl. Acad. Sci. 102, 6320–6325 (2005).

65. A. M. Bryan et al., Cholesterol and sphingomyelin are critical for Fcgamma receptor-mediated phagocytosis of Cryptococcus neoformans by macrophages. J Biol Chem 297, 101411 (2021).

66. S. Nazemidashtarjandi, V. M. Sharma, V. Puri, A. M. Farnoud, M. M. Burdick, Lipid Composition of the Cell Membrane Outer Leaflet Regulates Endocytosis of Nanomaterials through Alterations in Scavenger Receptor Activity. ACS Nano 16, 2233–2248 (2022).

67. V. Kainu, M. Hermansson, P. Somerharju, Electrospray ionization mass spectrometry and exogenous heavy isotope-labeled lipid species provide detailed information on aminophospholipid acyl chain remodeling. The Journal of biological chemistry 283, 3676–3687 (2008).

68. M. Refaei, R. Leventis, J. R. Silvius, Assessment of the roles of ordered lipid microdomains in post-endocytic trafficking of glycosyl-phosphatidylinositol-anchored proteins in mammalian fibroblasts. Traffic 12, 1012–1024 (2011).

69. B. B. Machta, S. Papanikolaou, J. P. Sethna, S. L. Veatch, Minimal model of plasma membrane heterogeneity requires coupling cortical actin to criticality. Biophys J 100, 1668–1677 (2011).

70. H. T. Cheng Megha, E. London, Preparation and properties of asymmetric vesicles that mimic cell membranes: effect upon lipid raft formation and transmembrane helix orientation. J Biol Chem 284, 6079–6092 (2009).

71. Q. Lin, E. London, Preparation of artificial plasma membrane mimicking vesicles with lipid asymmetry. PLoS One 9, e87903 (2014).

72. M. Son, E. London, The dependence of lipid asymmetry upon phosphatidylcholine acyl chain structure. J Lipid Res 54, 223–231 (2013).

73. V. T. Armstrong, M. R. Brzustowicz, S. R. Wassall, L. J. Jenski, W. Stillwell, Rapid flip-flop in polyunsaturated (docosahexaenoate) phospholipid membranes. Archives of Biochemistry and Biophysics 414, 74–82 (2003).

74. N. Bag, E. London, D. A. Holowka, B. A. Baird, Transbilayer Coupling of Lipids in Cells Investigated by Imaging Fluorescence Correlation Spectroscopy. J Phys Chem B 126, 2325–2336 (2022).

75. T. S. van Zanten et al., Direct mapping of nanoscale compositional connectivity on intact cell membranes. Proc Natl Acad Sci U S A 107, 15437–15442 (2010).

76. R. M. Young, D. Holowka, B. Baird, A lipid raft environment enhances Lyn kinase activity by protecting the active site tyrosine from dephosphorylation. J Biol Chem 278, 20746–20752 (2003).

77. D. Holowka et al., Lipid segregation and IgE receptor signaling: a decade of progress. Biochim Biophys Acta 1746, 252–259 (2005).

78. Y. Wang et al., Alpha-linolenic acid inhibits IgE-mediated anaphylaxis by inhibiting Lyn kinase and suppressing mast cell activation. Int Immunopharmacol 103, 108449 (2022).

