## Supporting Information for "Lipid Driven Inter-leaflet Coupling of Plasma Membrane Order Regulates FcεRI Signaling in Mast Cells"

**SUPPLEMENTARY FIGURES**

**Figure S1**

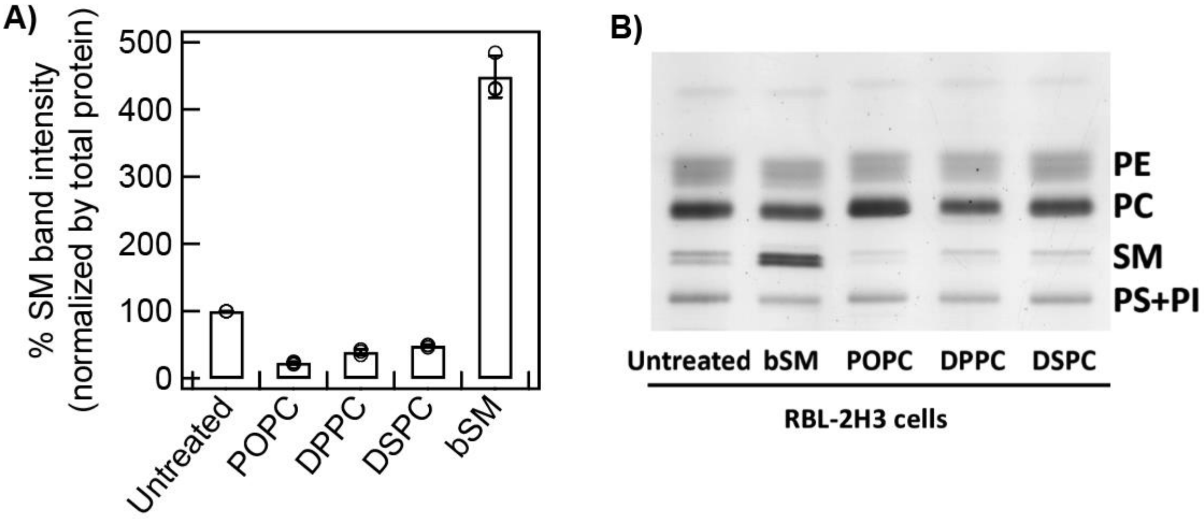


Figure S1. LEX efficiency in RBL cells is evaluated with SM loss in HP-TLC. A) %SM band intensity of the cells after lipid exchange with the specified lipids (POPC, DPPC, DSPC, and bSM) compared to treated RBL cells. Three independent HP-TLC experiments were done. The error bars represent standard deviation. B) A representative HP-TLC image is shown.

**Figure S2**

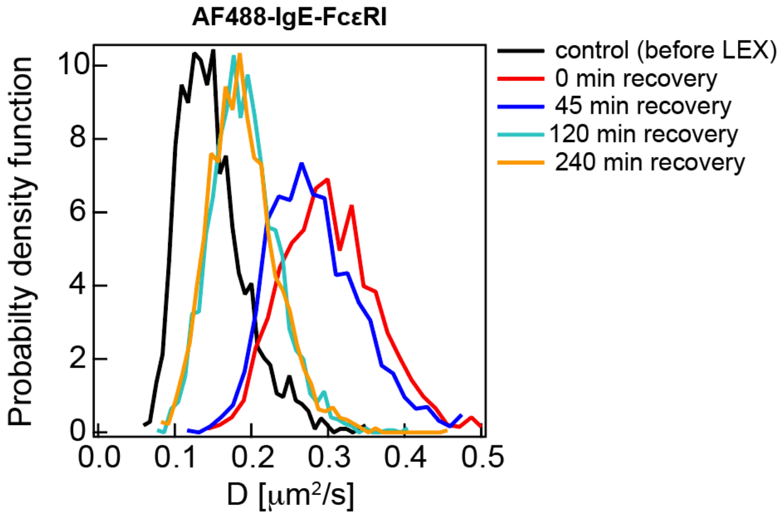


**Fig S2.** The diffusion coefficients (*D*) values of AF488-IgE-FcεRI recover to close to near control 120 min after LEX. The probability distribution functions of the D values are shown. The Number of cells at each time point = 5.

**Figure S3**

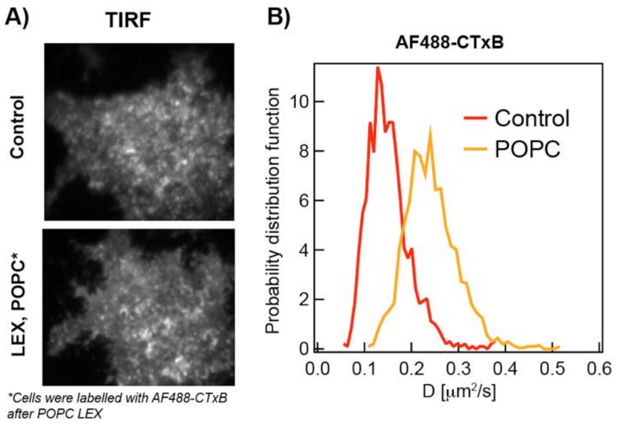


**Fig. S3:** POPC-LEX increases diffusion of AF488-CTxB-GM1 in outer leaflet. A) TIRF images of AF488-CTxB labelled unexchanged and POPC-LEX cells. B) The probability distribution functions of diffusion coefficients of AF488-CTxB-GM1 in unexchanged and POPC-LEX cells. Number of cells = 5 for each condition.

**Figure S4**

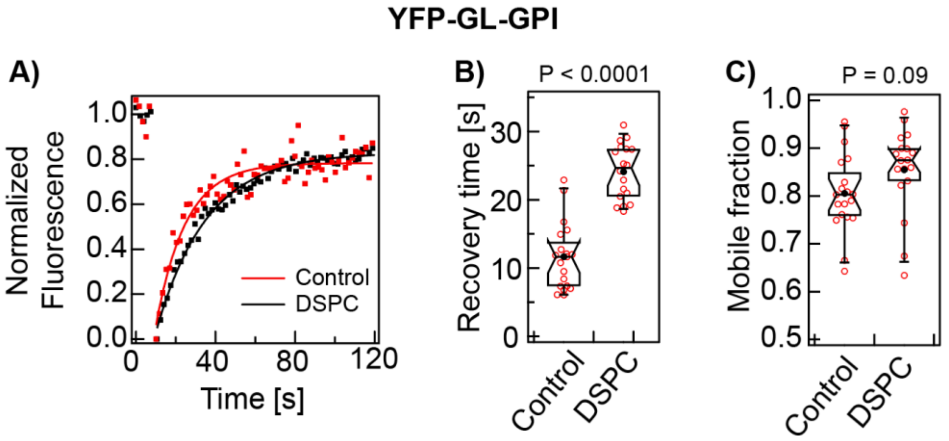


**Fig. S4:** DSPC-LEX does not induce gel-like regions in the outer leaflet as sensed by YFP-GL-GPI A) Representative recovery curve of individual cells. B) Recovery time, and C) mobile fraction of many cells.

**Figure S5**

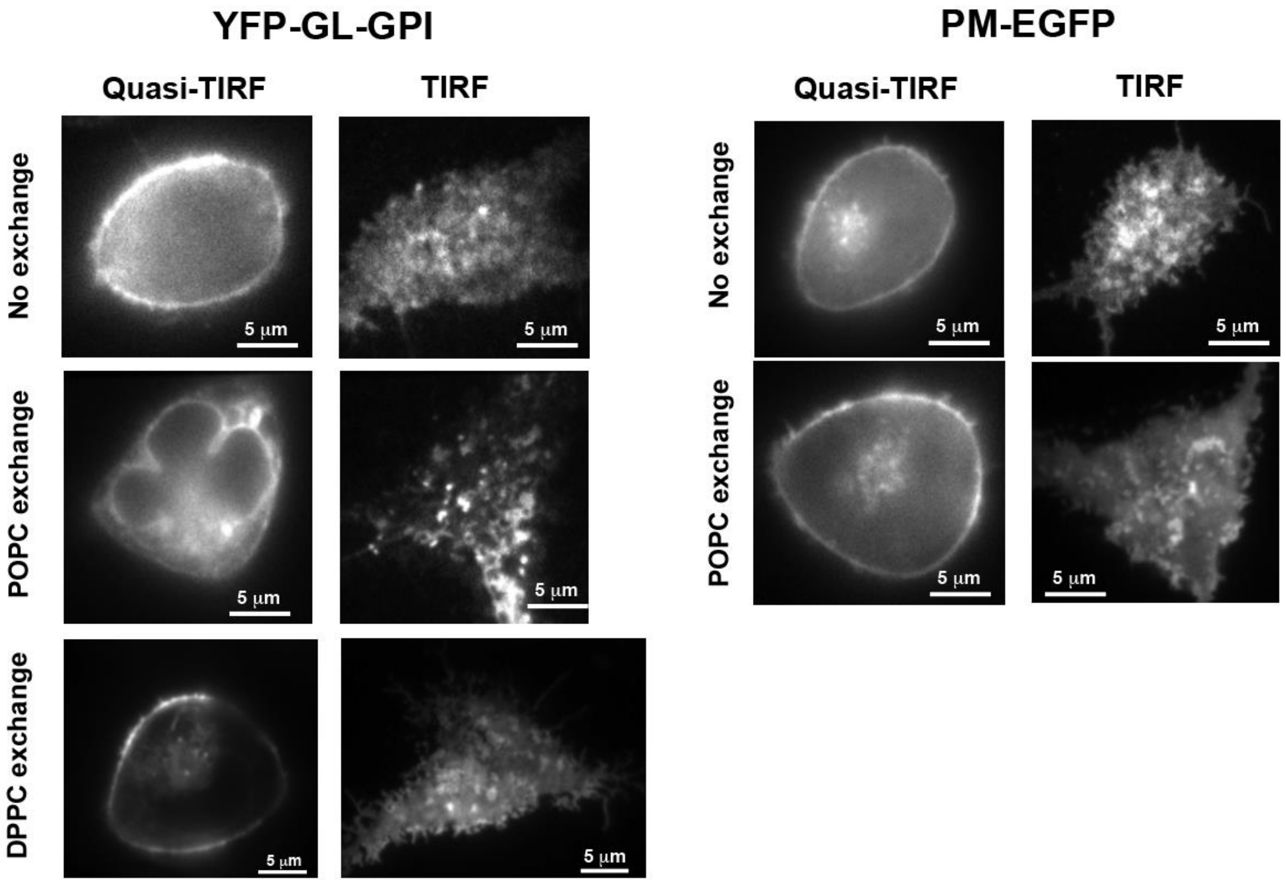


**Figure S5.** YFP-GL-GPI is internalized with POPC-LEX but not with DPPC-LEX. Left, quasi-TIRFM and TIRFM images of YFP-GL-GPI in unexchanged, POPC-LEX, and DPPC-LEX cells. Right, Quasi-TIRFM and TIRFM images of PM-EGFP in unexchanged and POPC-LEX cells.

### SUPPLEMENTARY MOVIE

**Movie S1:** Time-lapse TIRFM movie of stimulated YFP-Syk recruitment in a representative wildtype RBL cell. The movie is recorded at 1 min time resolution.
